# Potent activation of SARM1 by NMN analogue VMN underlies vacor neurotoxicity

**DOI:** 10.1101/2020.09.18.304261

**Authors:** Andrea Loreto, Carlo Angeletti, Weixi Gu, Andrew Osborne, Bart Nieuwenhuis, Jonathan Gilley, Peter Arthur-Farraj, Elisa Merlini, Adolfo Amici, Zhenyao Luo, Lauren Hartley-Tassell, Thomas Ve, Laura M. Desrochers, Qi Wang, Bostjan Kobe, Giuseppe Orsomando, Michael P. Coleman

**Affiliations:** John van Geest Centre for Brain Repair, Department of Clinical Neurosciences, University of Cambridge, Forvie Site, Robinson Way, CB2 0PY, Cambridge, UK; Department of Clinical Sciences (DISCO), Section of Biochemistry, Polytechnic University of Marche, Via Ranieri 67, Ancona 60131, Italy; School of Chemistry and Molecular Biosciences, Institute for Molecular Bioscience and Australian Infectious Diseases Research Centre, University of Queensland, QLD 4072, Australia; Institute for Glycomics, Griffith University, Southport, Queensland, 4215, Australia; Neuroscience, BioPharmaceuticals R&D, AstraZeneca, Waltham, MA, USA; Ikarovec Ltd, Norwich Innovation Centre, Norwich, UK; Cambridge Institute for Medical Research, University of Cambridge, Cambridge, United Kingdom; Vertex Pharmaceuticals, 50 Northern Ave, Boston, MA 02210, USA; Kymera Therapeutics, 200 Arsenal Yards Blvd, Watertown, MA 02472, USA

## Abstract

Axon loss underlies symptom onset and progression in many neurodegenerative disorders. Axon degeneration in injury and disease is promoted by activation of the nicotinamide adenine dinucleotide (NAD)-consuming enzyme SARM1 (sterile alpha and TIR motif-containing protein 1). Here, we report vacor mononucleotide (VMN), a metabolite of the pesticide and neurotoxin vacor, as the most potent yet SARM1 activator. Removal of SARM1 shows complete rescue from vacor-induced neuron and axon death *in vitro* and *in vivo*. We present the crystal structure of VMN bound to the *Drosophila* SARM1 regulatory armadillo-repeat domain, thus facilitating drug development to prevent SARM1 activation in human disease. This study indicates the likely mechanism of action of vacor as a pesticide and lethal neurotoxin in humans, provides important new tools for drug discovery, and further demonstrates that SARM1 removal can permanently block programmed axon death specifically induced by toxicity as well as genetic mutation.

## Introduction

Sterile alpha and TIR motif-containing protein 1 (SARM1) plays a central, pro-degenerative role in programmed axon death (including Wallerian degeneration) (Osterloh et al., 2012). This axon degeneration pathway is activated in a number of neurodegenerative contexts, including in human disease (Coleman and Höke, 2020; Gilley et al., 2021; Huppke et al., 2019; Lukacs et al., 2019). SARM1 has a critical nicotinamide adenine dinucleotide (NAD) cleavage (NADase) activity, which is activated when its upstream regulator and axon survival factor nicotinamide mononucleotide adenylyltransferase 2 (NMNAT2) is depleted or inactive (Essuman et al., 2017; Gilley and Coleman, 2010; Gilley et al., 2015; Sasaki et al., 2016, 2020a). NMNAT2 loss causes accumulation of its substrate nicotinamide mononucleotide (NMN), which promotes axon degeneration (Di Stefano et al., 2015, 2017; Loreto et al., 2015, 2020). NMN is now known to activate SARM1 NADase activity (Zhao et al., 2019). Given that *Sarm1* deletion confers robust axon protection, and even lifelong protection against lethality caused by NMNAT2 deficiency (Gilley et al., 2017), SARM1 has become a very attractive therapeutic target to prevent neurodegeneration. Understanding how SARM1 is activated by small molecule regulators will help develop ways to block its activation.

Here, we have investigated whether vacor, a disused pesticide and powerful neurotoxin associated with human peripheral and central nervous system disorders (Gallanosa et al., 1981; LeWitt, 1980) and axon degeneration in rats (Watson and Griffin, 1987), causes activation of programmed axon death. Vacor is metabolised to vacor mononucleotide (VMN) and vacor adenine dinucleotide (VAD) through a two-step conversion by nicotinamide phosphoribosyltransferase (NAMPT) and NMNAT2 (Fig. S1A). Considering VAD inhibits SARM1 regulator NMNAT2 (Buonvicino et al., 2018), we reasoned that vacor may induce SARM1-dependent axon death. We demonstrate that vacor neurotoxicity is entirely mediated by SARM1 such that *Sarm1*^-/-^ neurons and their axons are completely resistant. We unexpectedly find that vacor metabolite VMN directly binds to and activates SARM1, causing neuronal death. This study elucidates the mechanism of action of vacor likely underlying its lethal neurotoxic effect in humans, and provides important new tools for drug discovery. To our knowledge, VMN is the most potent activator of SARM1 reported so far, and our structural data reveal essential information on SARM1 regulation which will aid drug development to block SARM1 activation.

## Results

### Vacor causes SARM1-dependent neurite and cell death

Intracellular conversion of vacor into its metabolites has been suggested to contribute to its toxicity (Buonvicino et al., 2018). We first confirmed that VMN and VAD are generated in vacor-treated dorsal root ganglion (DRG) mouse neurons (Fig. S1B) and that these and a second neuron type, superior cervical ganglia (SCG) mouse neurons, exhibit rapid, dose-dependent neurite degeneration (Fig. 1A-D; Fig. S1C-F). Consistent with the proposed toxic role for vacor metabolites (Buonvicino et al., 2018), both nicotinamide, which competes with vacor as a preferred substrate for NAMPT, and the NAMPT inhibitor FK866 (Fig. S1A), delayed vacor-induced neurite degeneration (Fig. S1G-J). Such competition may explain why NAM was an effective treatment for patients with vacor poisoning (Gallanosa et al., 1981).

**Fig. 1.**
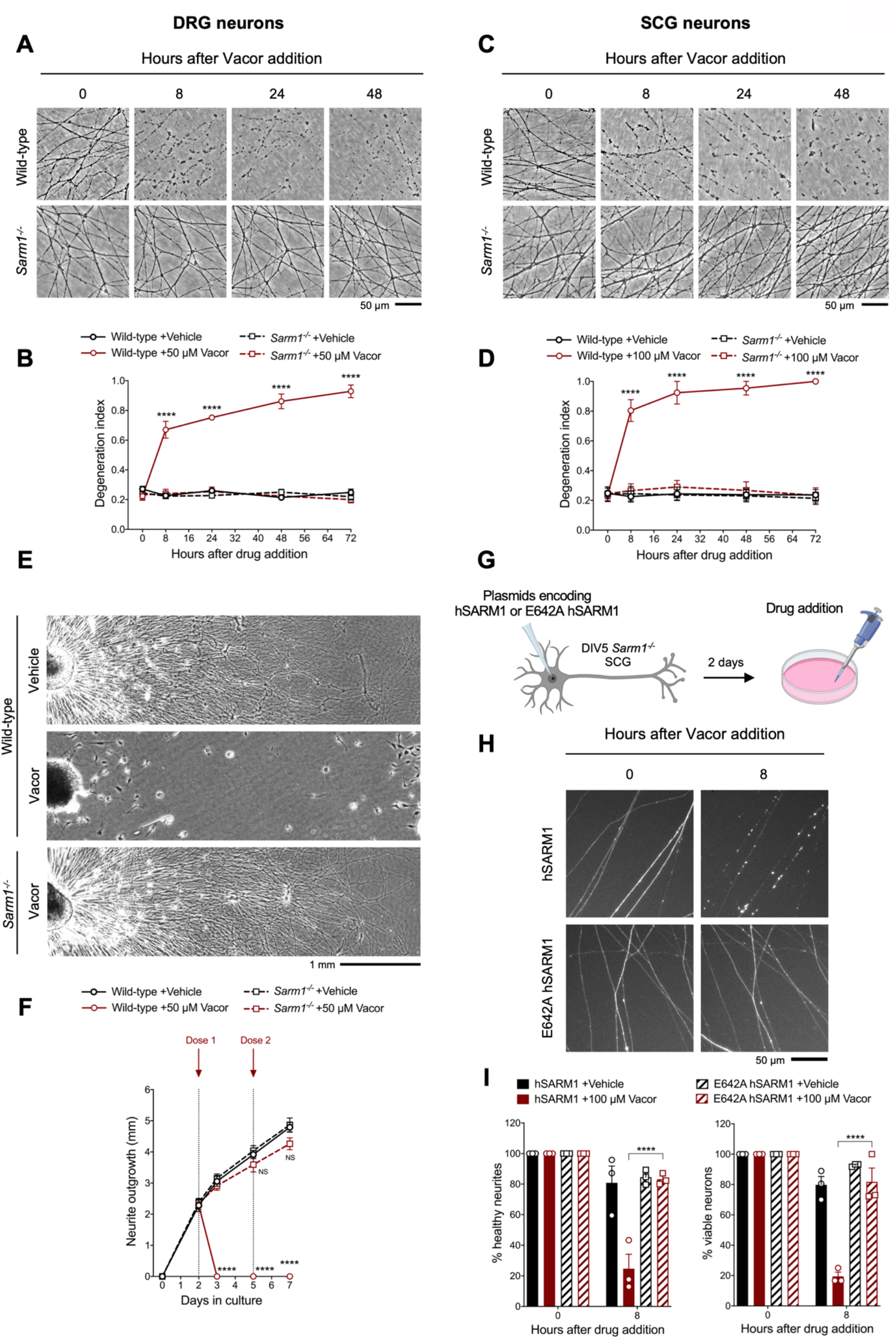
Vacor causes SARM1-dependent neurite and cell death. (See also Fig. S1-3). **(A)** Representative images of neurites from wild-type and *Sarm1*^*-/-*^ DRG (littermates) explant cultures treated with 50 µM vacor. **(B)** Quantification of the degeneration index in experiments described in (A) (mean ± SEM; n = 4; repeated measures three-way ANOVA followed by Tukey’s multiple comparison test; ****, p<0.0001; statistical significance shown relative to *Sarm1*^*-/-*^ +50 µM Vacor). **(C)** Representative images of neurites from wild-type and *Sarm1*^*-/-*^ SCG explant cultures treated with 100 µM vacor. **(D)** Quantification of the degeneration index in experiments described in (C) (mean ± SEM; n = 4; repeated measures three-way ANOVA followed by Tukey’s multiple comparison test; ****, p < 0.0001; statistical significance shown relative to *Sarm1*^*-/-*^+100 µM Vacor). **(E)** Representative images of neurite outgrowth at DIV7 from wild-type and *Sarm1*^*-/-*^ DRG explant cultures treated with 50 µM vacor or vehicle. Multiple doses of vacor or vehicle were added at DIV2 and DIV5. **(F)** Quantification of neurite outgrowth in (E) (mean ± SEM; n = 5; repeated measures three-way ANOVA followed by Tukey’s multiple comparison test; ****, p < 0.0001; NS, not-significant; statistical significance shown relative to Wild-type +Vehicle). **(G)** Schematic representation of the experimental design for (H) (‘Created with BioRender’). **(H)** Representative images of neurites from *Sarm1*^*-/-*^ SCG dissociated neurons co-injected with plasmids encoding wild-type or E642A hSARM1 and DsRed (to label neurites) and treated with 100 µM vacor. **(I)** Quantification of healthy neurites and viable neurons in experiments in (H) is shown as a percentage relative to 0 hr (time of drug addition) (mean ± SEM; n = 3; repeated measures three-way ANOVA followed by Tukey’s multiple comparison test; ****, p < 0.0001).

As hypothesised, vacor failed to induce degeneration of *Sarm1*^*-/-*^ DRG and SCG neurites (Fig. 1A-D). This protection was extremely strong, with neurites surviving indefinitely even after multiple vacor doses (Fig. S2A,B). *Sarm1*^*-/-*^ neurites were not only protected from degeneration, but they also continued to grow normally, even with repeated dosing (Fig. 1E,F). This mirrors the permanent rescue and continued growth previously reported in *Nmnat2* null axons in the absence of SARM1 (Gilley et al., 2015, 2017), suggesting complete efficacy in both toxic and inherited types of neuropathy. It also shows that vacor neurotoxicity is predominantly SARM1-dependent.

SARM1 levels remained relatively stable following vacor treatment (Fig. S2C-F). We therefore tested if SARM1 NADase activity is the critical function required for vacor toxicity. Exogenous expression of wild-type human SARM1 (hSARM1) in *Sarm1*^*-/-*^ SCG neurons restored vacor sensitivity, whereas expression of E642A hSARM1, which lacks NADase activity, did not (Fig. 1G-I; Fig. S2G,H), similar to findings in axotomy (Essuman et al., 2017). Interestingly, unlike axotomy, activation of the pathway by vacor also caused SARM1-dependent cell death (Fig. 1I), similar to that previously reported with constitutively active SARM1 (Gerdts et al., 2013, 2015). To explore this further, we cultured DRG neurons in microfluidic chambers, to allow independent manipulation of the neurite and soma compartments. Addition of vacor to either compartment directly activated local death of cell bodies and/or neurites. However, while vacor-induced cell death causes eventual secondary neurite degeneration, the reverse was not true: no cell loss was observed after local induction of distal neurite degeneration by vacor (Fig. S3A,B).

### *Sarm1* deletion confers functional and morphological protection of neurons against vacor toxicity *in vivo*

We next sought to validate our findings *in vivo*. To reduce the impact on animal welfare of systemic administration of a rodenticide in mice, we opted for intravitreal (IVT) injection (Fig. 2A) of vacor. Electroretinogram (ERG) recordings showed that vacor caused a global loss of photoreceptor neuronal activity, and subsequent loss of bipolar and retinal ganglion cell (RGC) responses, which were fully rescued by SARM1 deficiency (Fig. 2B,C). Vacor administration also caused inner retina neuronal death, as revealed by a significant reduction in RGC number. Again, this was completely prevented by SARM1 deficiency (Fig. 2D,E). DMSO had no adverse effect on retinal cell survival or function compared to PBS-injected eyes (Fig. S4A,B). These data demonstrate that the absence of SARM1 confers functional and morphological protection of neurons against vacor toxicity *in vivo*.

**Fig. 2.**
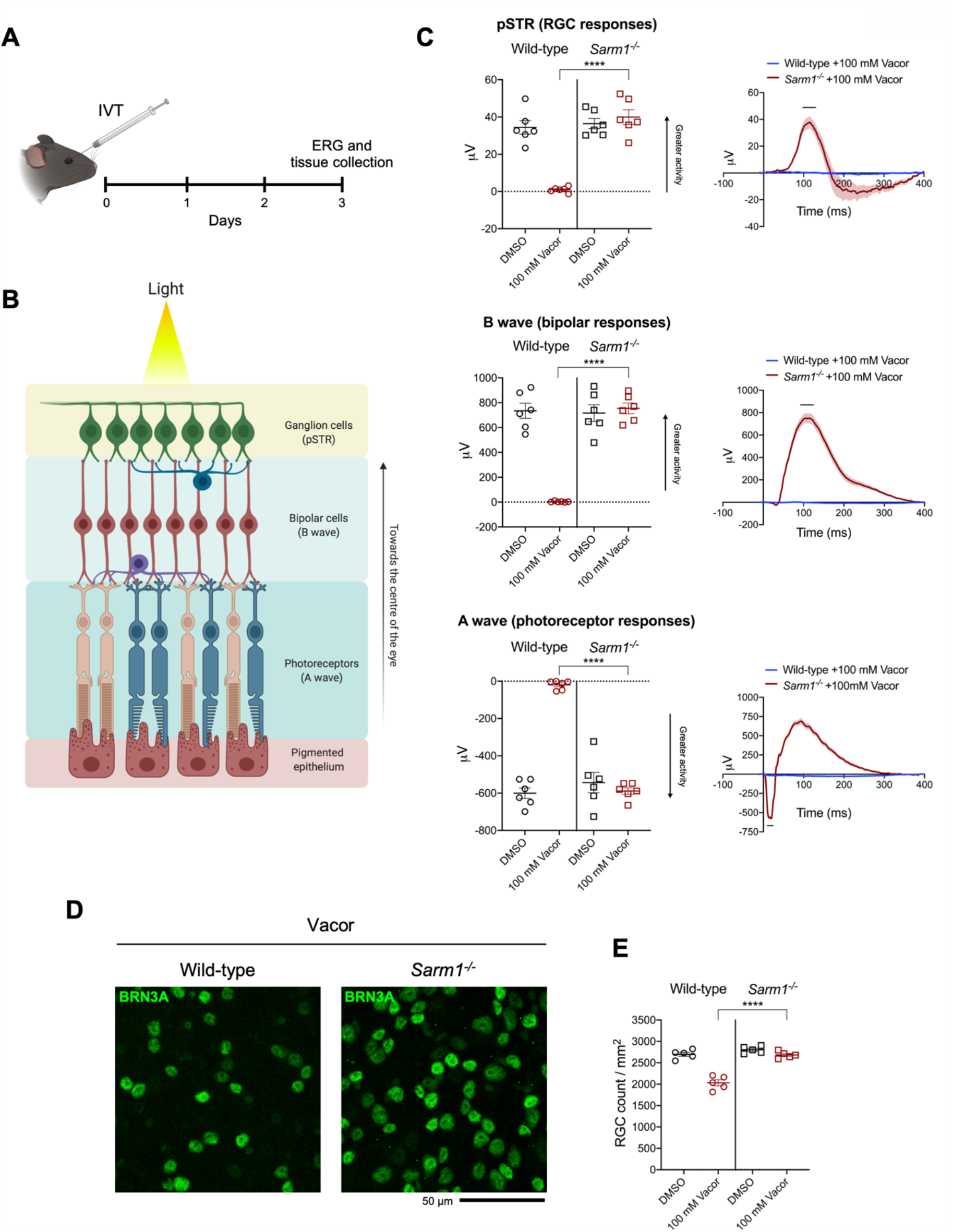
*Sarm1* deletion confers functional and morphological protection of neurons against vacor toxicity *in vivo*. (See also Fig. S4). **(A)** Schematic representation of the experimental design for (C,D) (‘Created with BioRender’). **(B)** Graphic illustration of the different retinal layers (‘Created with BioRender’). **(C)** Quantification of the ERG responses (pSTR, B wave and A wave) from wild-type and *Sarm1*^*-/-*^ mice injected with 100 mM vacor or DMSO (vehicle) (mean ± SEM; n = 6; two-way ANOVA followed by Tukey’s multiple comparison test; ****, p < 0.0001). **(D)** Representative images of RGC from wild-type and *Sarm1*^*-/-*^ mice injected with 100 mM vacor or DMSO (vehicle). **(E)** Quantification of RGC numbers from wild-type and *Sarm1*^*-/-*^ mice injected with 100 mM vacor or DMSO (vehicle) (mean ± SEM; n = 5; repeated measures three-way ANOVA followed by Tukey’s multiple comparison test; ****, p<0.0001).

### Vacor treatment leads to SARM1 activation

Activation of SARM1 depletes NAD and generates cADPR in neurons (Essuman et al., 2017; Gerdts et al., 2015; Sasaki et al., 2016, 2020a). Based on the previously reported inhibition of NMNAT2 by VAD (Buonvicino et al., 2018), we initially hypothesised that this causes a rise in NMN, the physiological NMNAT2 substrate and a known activator of SARM1 (Di Stefano et al., 2015, 2017; Loreto et al., 2015; Zhao et al., 2019), thereby stimulating SARM1 activity to trigger vacor-dependent axon death. Surprisingly, while SARM1 activation was confirmed by a drastic SARM1-dependent decline in NAD and increase in cADPR following vacor administration, NMN did not rise and even fell slightly (Fig. 3A-D). Furthermore, NMN and NAD levels and their ratio to one another were not altered in our vacor-treated *Sarm1*^*-/-*^ neurons (Fig. 3A,B), suggesting that the reported inhibition of NAMPT and NMNAT2 by vacor/VAD (Buonvicino et al., 2018) does not occur in this context at this drug concentration. Crucially, NMN deamidase, an enzyme that strongly preserves axotomised axons by preventing NMN accumulation (Fig. 3E; Fig. S5A) (Di Stefano et al., 2015, 2017; Loreto et al., 2015), was also unable to protect against vacor toxicity (Fig. 3E,F), even though recombinant NMN deamidase retains its ability to convert NMN to NaMN in the presence of vacor, VMN and VAD (Fig. S5B). WLD^S^, another enzyme that limits NMN accumulation (Di Stefano et al., 2015), also failed to rescue neurons against vacor neurotoxicity (Fig. S5C,D). These data suggest that, in this specific context at least, NMN is not responsible for SARM1 activation.

**Fig. 3.**
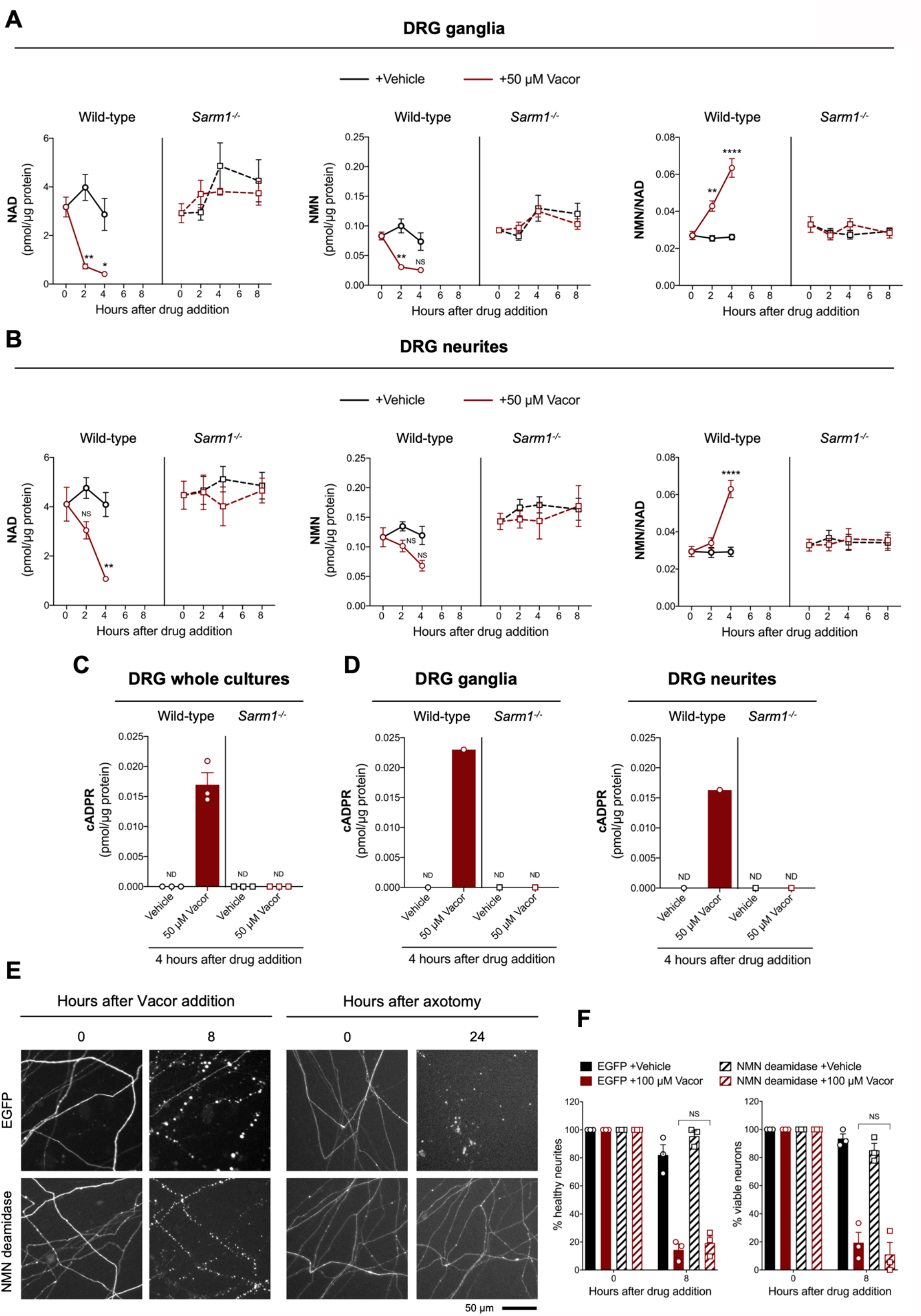
Vacor treatment leads to SARM1 activation. (See also Fig. S5). **(A,B)** NMN and NAD levels and NMN/NAD ratio in ganglia (A) and neurite (B) fractions from wild-type and *Sarm1*^-/-^ DRG explant cultures at the indicated time points after 50 µM vacor or vehicle treatment (mean ± SEM; n = 4; three-way ANOVA followed by Tukey’s multiple comparison test; ****, p < 0.0001; **, p < 0.01; *, p < 0.05; NS, not-significant). **(C)** cADPR levels in wild-type and *Sarm1*^-/-^ DRG whole explant cultures (neurites and cell bodies) 4 hr after 50 µM vacor or vehicle treatment. cADPR levels were consistently above the detection limit (∼1 fmol/µg protein) only in wild-type DRG explant cultures treated with vacor (mean ± SEM; n = 3; ND, not-detectable). **(D)** A single analysis of cADPR levels in ganglia and neurite fractions from wild-type and *Sarm1*^-/-^ DRG explant cultures 4 hr after 50 µM vacor or vehicle treatment (n = 1). **(E)** Representative images of neurites from *Sarm1*^*-/-*^ SCG dissociated neurons co-injected with plasmids encoding hSARM1, EGFP or EGFP-NMN deamidase and DsRed (to label neurites) and treated with 100 µM vacor. As an experimental control, *Sarm1*^*-/-*^ SCG dissociated neurons injected with the same injection mixtures were axotomised. As expected, neurites expressing NMN deamidase were still intact 24 hr after axotomy. **(F)** Quantification of healthy neurites and viable neurons in experiments in (A) is shown as a percentage relative to 0 hr (time of drug addition) (mean ± SEM; n = 3; repeated measures three-way ANOVA followed by Tukey’s multiple comparison test; NS, not-significant).

### Vacor metabolite VMN potently activates SARM1 through direct binding to the ARM domain

What did accumulate in vacor-treated neurons was the vacor metabolite VMN (Fig. S1B) and, given its structural similarity to the endogenous SARM1 activator NMN (Fig. S6A), we hypothesised that VMN might instead directly activate SARM1, leading to cell and axon death. Crucially, we found that VMN potently activates the NADase activity of recombinant hSARM1, even more so than NMN, having a lower K_a_ and resulting in a greater induction (Fig. 4A; Fig. S6B). Conversely, VAD only had a weak inhibitory effect on hSARM1 activity (Fig. S6C). Intriguingly, recombinant hSARM1 NADase activity dropped at higher VMN concentrations (Fig. 4A). Notwithstanding the doses of vacor used in this study resulting in VMN levels in neurons within the activation range of SARM1, this inhibitory action could reveal critical information on how SARM1 activity is regulated. Analysis via best fitting (see equation in Methods) suggests that our data are compatible with two distinct binding sites of VMN on SARM1 and led us to establish binding constants for activation (K_a_) and inhibition (K_i_) (Fig. 4A).

**Fig. 4.**
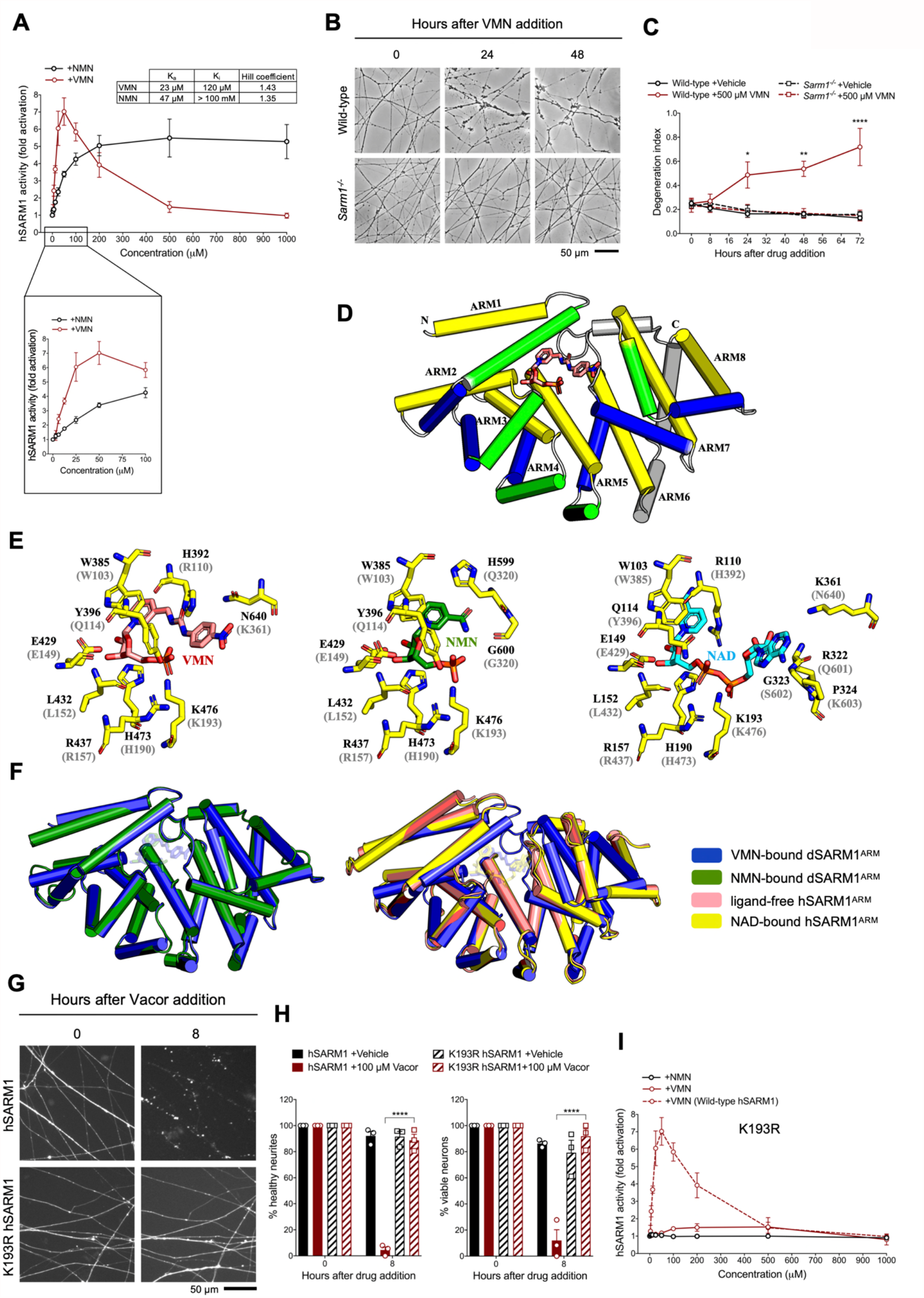
VMN potently activates SARM1 through direct binding to the ARM domain. (See also Fig. S6-9). **(A)** Fold change of NADase activity of purified, recombinant hSARM1 in the presence of NMN and VMN (mean ± SEM; n = 3). hSARM1 average basal activity is 18.12 ± 3.02 milliU/mg (fold activation = 1). Rates are relative to controls measured with 250 µM NAD alone. Both NMN and VMN, once added to each reaction mixture, were not consumed during incubation. Experimental data were fitted to the modified Michaelis-Menten equation in Methods to calculate the kinetic parameters shown in the attached table. Hill coefficients for NMN and VMN indicate positive cooperativity in binding in both cases. Best fitting also revealed a *K*_m_ for NAD of 70 µM. **(B)** Representative images of neurites from wild-type and *Sarm1*^*-/-*^ DRG (littermates) explant cultures treated with 500 µM VMN. **(C)** Quantification of the degeneration index in experiments described in (B) (mean ± SEM; n = 3; repeated measures three-way ANOVA followed by Tukey’s multiple comparison test; ****, p < 0.0001; **, p < 0.01; *, p < 0.05; statistical significance shown relative to *Sarm1*^*-/-*^ +500 µM VMN). **(D)** Crystal structure of VMN-bound dSARM1^ARM^. dSARM1^ARM^ contains 8 armadillo motifs (ARM1-8). Except for ARM1 (containing only the H3 helix) and ARM7(containing only H2 and H3 helices), other motifs consist of H1, H2 and H3 helices, coloured green, blue and yellow, respectively. The unusual ARM6, which makes a sharp turn at its C-terminus, is coloured grey. Nitrogen, oxygen and phosphorous are coloured blue, red and orange, respectively, in the VMN molecule. **(E)** Stick representation of the interaction of dSARM1^ARM^ with VMN (pink; left) and NMN (green; middle), and hSARM1^ARM^ with NAD (PDB: 7CM6; cyan; right). The corresponding residues in *Drosophila* or human SARM1^ARM^ are shown in parentheses. Nitrogen, oxygen and phosphorous are coloured blue, red and orange. **(F)** Structural comparison of the ARM domains bound to different ligands. The panel on the left shows the structural superposition of VMN and NMN-bound dSARM1^ARM^ (PDB: 7LCZ; RMSD is 0.4 Å over 304 Cα atoms). Panel on the right shows the superposition of VMN-bound dSARM1^ARM^, unbound (PDB: 7CM5) and NAD-bound hSARM1^ARM^ (PDB: 7CM6) using the function cealign in PyMol (RMSD between ligand-free and VMN-bound structures is 2.7 Å over 288 Cα atoms; RMSD between NAD-bound and VMN-bound structures is 2.9 Å over 288 Cα atoms). **(G)** Representative images of neurites from *Sarm1*^*-/-*^ SCG dissociated neurons co-injected with plasmids encoding wild-type or K193R hSARM1 and DsRed (to label neurites) and treated with 100 µM vacor. **(H)** Quantification of healthy neurites and viable neurons in experiments in (E) is shown as a percentage relative to 0 hr (time of drug addition) (mean ± SEM; n = 3; repeated measures three-way ANOVA followed by Tukey’s multiple comparison test; ****, p < 0.0001). **(I)** Fold change of NADase activity of purified, recombinant K193R hSARM1 in the presence of NMN and VMN (wild-type hSARM1 + VMN is also shown for comparison) (mean ± SEM; n = 2-3). K193R hSARM1 average basal activity is 17.75 ± 2.47 milliU/mg (fold activation = 1).

While NMN activates SARM1 (Zhao et al., 2019) and promotes programmed axon death when NMNAT2 is depleted (Di Stefano et al., 2015, 2017; Loreto et al., 2015, 2020), exogenous NMN does not induce degeneration of uninjured neurites (Di Stefano et al., 2015), probably because it is rapidly converted to NAD when NMNAT2 is present. However, we now show that exogenous application of its analogue, VMN, does induce SARM1-dependent death of uninjured DRG neurites (Fig. 4B,C). This difference likely reflects a combination of VMN being both a more potent activator of SARM1 and it accumulating more because it is not efficiently metabolised by NMNAT2 (Buonvicino et al., 2018). VMN-dependent activation of SARM1 being the effector of vacor toxicity is also supported by several other findings. First, unlike NMN, VMN is not a substrate of NMN deamidase (Fig. S6D), thus explaining its inability to protect against vacor neurotoxicity (Fig. 3E,F). In addition, VMN is not a substrate of the nuclear isoform NMNAT1 and is a poor substrate of mitochondrial NMNAT3 (Buonvicino et al., 2018), so VMN accumulation in cell bodies (Fig. S6E) and subsequent SARM1 activation, as detected by a rise of cADPR in this compartment (Fig. 3D), provides a clear rationale for why vacor administration also causes cell death.

SARM1 armadillo-repeat (ARM) domain is believed to maintain SARM1 in an inhibited state (Figley et al., 2021; Gerdts et al., 2013; Jiang et al., 2020). SARM1 is activated by NMN binding directly to the ARM domain (Figley et al., 2021; Gu et al., 2021). We therefore hypothesised that VMN could directly interact with the ARM domain to release this inhibition and activate SARM1, causing neuronal death. Because we were unable to produce hSARM1^ARM^ recombinantly, we used the ARM domain of *Drosophila* SARM1 (dSARM1^ARM^, residues 315-678), produced in *E. coli*, to investigate if VMN interacts directly with dSARM1^ARM^. ITC (isothermal titration calorimetry) analysis showed that VMN bound to dSARM1^ARM^ with a K_d_ value of 2.83 ± 0.16 μM (1:1 molar ratio). To further characterise this interaction, we determined the crystal structure of the VMN-bound dSARM1^ARM^ (1.69 Å resolution). dSARM1^ARM^ contains 8 tandem armadillo repeats (ARM1-8) stacking into an unusually compact right-handed superhelix (Fig. 4D; Table S1). The bound VMN molecule sits in the central groove of dSARM1^ARM^, which is mainly lined by the H3 helices from ARM1-5 (Fig. 4D; Fig. S7B). The interaction is mediated by two pi-stacking interactions between the indole of W385 and the pyridine moiety of VMN, and between the imidazole ring of H392 and the nitrobenzene moiety of VMN (Fig. 4E; Fig. S8A,B); and 8 hydrogen bonds, with a buried surface area of ∼910 Å^2^ (Fig. 4E; Fig. S8A,B). The hydroxyl group from the side chain of Y396 and the amino groups from the side chains of the conserved R437 and K476 form hydrogen bonds with the phosphate group of VMN; the amino group from the side chain of N640 forms a hydrogen bond with the nitro group of VMN; and the imidazole rings of H392 and H473 form hydrogen bonds with the urea and ribose of VMN, respectively (Fig. 4E; Fig. S8A,B).

Our data show that VMN has over two-fold higher affinity for dSARM1^ARM^ than NMN (K_d_ = 6.39 ± 0.04 μM), whose own interaction with the ARM domain was recently reported (Fig. S7A) (Figley et al., 2021). VMN-bound dSARM1^ARM^ adopts a similar conformation to NMN-bound dSARM1^ARM^ (Fig. 4F). Comparison of the two structures reveals that the NMN moiety of VMN and NMN share the same binding mode, with the key residues involved in NMN binding interacting with the analogous portions of the VMN molecule (Fig. 4E). Importantly, in the NMN-bound structure, NMN forms a hydrogen bond with the residues H599 and G600 (loop connecting ARM6 and 7), while in the VMN-bound structure, due to the extra portion of VMN (urea and nitrobenzene groups) extending to the region between the N- and C-termini of the protein, two different hydrogen bonds are formed between VMN and H392 (ARM2 H1) at the N-terminus, and N640 (ARM8 H1) at the C-terminus. Furthermore, the pi-stacking interaction between H392 and the nitrobenzene of VMN is specific to this compound. These different and extra interactions VMN forms could explain why VMN induces SARM1 activation more efficiently, compared to NMN. The compaction seen in both NMN and VMN-bound dSARM1^ARM^ structures likely represents the conformation of the ARM domain in active SARM1, based on the comparison with the more open cryo-EM (electron microscopy) structures of ligand-free and NAD-bound hSARM1, representing the inactive state of the protein (Bratkowski et al., 2020; Figley et al., 2021; Sporny et al., 2020) (Fig. 4F). In the NAD-bound structure, the portion of NAD equivalent to NMN and VMN interacts with the protein through similar interactions, but the adenine group of NAD extends further down to the side of the region between the termini. This would cause steric clashes with *Drosophila* N640 (predicted to correspond to human K361), thus preventing NAD from inducing a more compact conformation in the ARM domain (Fig. S7C), keeping the protein in the inactive state.

In summary, our data suggest that VMN activates SARM1 with a mechanism similar to NMN but with some important differences, inducing a compaction of the ARM domain, destabilising its interfaces with the neighbouring ARM, SAM and TIR domains, and eventually permitting the TIR domains to self-associate and hydrolyse NAD.

To validate the interactions observed in the crystal structure, we performed mutagenesis studies, focusing on the interactions associated with W385 (human W103), R437 (human R157) and K476 (human K193), because these residues are highly conserved and specifically interact with VMN through their side chains. We introduced the corresponding W103A, R157A and K193R mutations into human SARM1 and exogenously expressed them in *Sarm1*^*-/-*^ SCG mouse neurons (Fig. S9C). Unlike wild-type hSARM1, these mutants failed to restore neurite degeneration and cell death after vacor administration (Fig. 4G,H; Fig. S9A,B). We picked K193R hSARM1 for further biochemical characterisation, as it is known to cause loss-of-function, based on a previous study (Geisler et al., 2019). Crucially, neither VMN nor NMN could activate the NADase activity of recombinant K193R hSARM1 (Fig. 4I).

## Discussion

Overall, our study provides clear evidence that direct SARM1 activation by the vacor metabolite VMN underlies vacor neurotoxicity. Crucially, the comparison between VMN and NMN-bound dSARM1^ARM^ structures and the differences from the ligand-free and NAD-bound hSARM1^ARM^ structures provide the explanation for the mechanism of how these ligands target the same allosteric site, but regulate SARM1 activity differently. The data not only provide support for a key physiological role of NMN in the regulation of SARM1 activity and axon degeneration, but given VMN’s greater potency than NMN, we anticipate that VMN and vacor will be important tools to support drug discovery in several ways. The identification of key residues in the ARM domain involved in the potent activation of SARM1 by VMN should facilitate rational drug design; further understanding why VMN activates SARM1 more potently than NMN could lead to the development of ligands with higher-affinity and selectivity to lock SARM1 in its inactive state. It will be important to expand also on VMN inhibition functions. Furthermore, vacor is currently the most effective chemical to directly activate SARM1 and cause neuronal death, eliminating the need for complex axotomy experiments or the use of drugs with non-specific actions. The IVT vacor model we have developed could also be used for relatively rapid validation of drugs targeting SARM1 *in vivo* in rodents, with both morphological and functional readouts. Our data also show that SARM1 activation is sufficient to drive degeneration of multiple types of retinal neuron in addition to photoreceptors (Sasaki et al., 2020b), making SARM1 a promising target to treat retinal degeneration.

Our data also further implicate programmed axon death in human disease. It appears that this degeneration pathway can be aberrantly activated in humans not only by genetic mutation of *NMNAT2* (Huppke et al., 2019; Lukacs et al., 2019), but also by SARM1 activation in severe neurotoxicity within hours of vacor ingestion (LeWitt, 1980). Although vacor use is banned, this study raises important questions of whether other pesticides or environmental chemicals in use today also activate programmed axon death. Despite the extreme toxicity of vacor, SARM1 deficiency completely rescues neurons and preserves their functionality. Together with the previously reported lifelong protection in a genetic model of a rare human axonopathy (Gilley et al., 2017; Lukacs et al., 2019), this further supports the therapeutic potential of blocking SARM1 to completely prevent programmed axon death. Lastly, as vacor causes pancreatic β-cell destruction and diabetes in humans (Gallanosa et al., 1981), these findings could also have broader implications, uncovering a role for SARM1 in the survival of insulin producing cells and aiding research on diabetes. In conclusion, our study identifies a powerful SARM1 activator, advancing our understanding of how SARM1 activity is regulated and providing key support to drug discovery targeting programmed axon death.

## Methods

All studies conformed to the institution’s ethical requirements in accordance with the 1986 Animals (Scientific Procedures) Act, and in accordance with the Association for Research in Vision and Ophthalmology’s Statement for the Use of Animals in Ophthalmic and Visual Research.

### Primary neuronal cultures

DRG ganglia were dissected from C57BL/6Babr, *Sarm1*^-/-^ and *Wld*^*S*^ E13.5-E14.5 mouse embryos and SCG ganglia were dissected from postnatal day 0-2 mouse pups. Littermates from *Sarm1*^+/-^ crosses were used when possible, as indicated in figure legends. Explants were cultured in 35 mm tissue culture dishes pre-coated with poly-L-lysine (20 µg/ml for 1 hr; Merck) and laminin (20 µg/ml for 1 hr; Merck) in Dulbecco’s Modified Eagle’s Medium (DMEM, Gibco) with 1% penicillin/streptomycin, 50 ng/ml 2.5S NGF (all Invitrogen) and 2% B27 (Gibco). 4 µM aphidicolin (Merck) was used to reduce proliferation and viability of small numbers of non-neuronal cells (Loreto and Gilley, 2020). For cultures of dissociated SCG neurons, *Sarm1*^-/-^ SCG ganglia were incubated in 0.025% trypsin (Merck) in PBS (without CaCl_2_ and MgCl_2_) (Merck) for 30 min, followed by incubation with 0.2% collagenase type II (Gibco) in PBS for 20 min. Ganglia were then gently dissociated using a pipette. Dissociated neurons were plated in a poly-L-lysine and laminin-coated area of ibidi μ-dishes (Thistle Scientific) for microinjection experiments. Dissociated cultures were maintained as explant cultures, except that B27 was replaced with 10% fetal bovine serum (Merck) and 2.5S NGF was lowered to 30 ng/ml. Culture media was replenished every 3 days. For most experiments, neurites were allowed to extend for 7 days before treatment.

### Drug treatments

For most experiments, DRG and SCG neurons were treated at day *in vitro* (DIV) 7 with vacor (Greyhound chromatography) or vehicle (H_2_O with 4% 1 N HCl), and VMN or vehicle (H_2_O) just prior to imaging (time 0 hr). When used, FK866 (kind gift of Prof Armando Genazzani, University of Novara) and NAM (Merck) were added at the same time as vacor. For neurite outgrowth and long-term survival assays, multiple doses of vacor or vehicle were added by replacing media with fresh media containing the drugs at the timepoints indicated in the figure. The drug concentrations used are indicated in the figures and figure legends. Vacor was dissolved in H_2_O with 4% 1 N HCl or DMSO; quantitation of the dissolved stock was performed spectrophotometrically (ɛ_340nm_ 17.8 mM^-1^cm^-1^).

### Acquisition of phase contrast images and quantification of neurite degeneration and outgrowth

Phase contrast images were acquired on a DMi8 upright fluorescence microscope (Leica microsystems) coupled to a monochrome digital camera (Hamamatsu C4742-95). The objectives used were NPLAN 5X/0.12 for neurite outgrowth assays and HCXPL 20X/0.40 CORR for neurite degeneration assays. Radial outgrowth was determined by taking the average of two measurements of representative neurite outgrowth for each ganglion at DIV2-3-5-7. Measurements were made from overlapping images of the total neurite outgrowth. For neurite degeneration assays, the degeneration index was determined using an ImageJ plugin (Sasaki et al., 2009). For each experiment, the average was calculated from three fields per condition; the total number of experiments is indicated in the figure legends.

### Microinjection and quantification of % of healthy neurites and viable neurons

DIV5 dissociated *Sarm1*^-/-^ SCG neurons were microinjected using a Zeiss Axiovert S100 microscope with an Eppendorf FemtoJet microinjector and Eppendorf TransferMan® micromanipulator. Plasmids were diluted in 0.5X PBS (without CaCl_2_and MgCl_2_) and filtered using a Spin-X filter (Costar). The mix was injected directly into the nuclei of SCG neurons using Eppendorf Femtotips (Gilley and Loreto, 2020). Injected plasmids were allowed to express for 2 days before vacor or vehicle treatment and axotomy. Plasmids were injected at the following concentrations: 2.5 (Fig. 1G-I) or 10 (Fig. 3E,F) ng/μl (untagged) wild-type hSARM1 and 2.5 ng/μl E642A hSARM1 (pCMV-Tag2 vector backbone), 10 ng/μl (C-terminal Flag-tagged) wild-type, K193R, R157A and W103A hSARM1 (pLVX-IRES-ZsGreen vector backbone) (Fig. 4G,H; Fig. S9A,B), 30 ng/μl EGFP-NMN deamidase (pEGFP-C1 vector backbone), 30 ng/μl pEGFP-C1, 40 (Fig. 1G-I; Fig. 4G,H; Fig. S9A,B) and 70 (Fig. 3E,F) ng/μl pDsRed2-N1. To check for expression of wild-type and E642A hSARM1 constructs (pCMV-Tag2 vector backbone) (Fig. S2H), wild-type, K193R, R157A and W103A hSARM1 (pLVX-IRES-ZsGreen vector backbone) (Fig. S9C) and pDsRed2-N1 were injected at 25 ng/μl; neurons were then fixed in 4% PFA (Merck) and immunostained with mouse monoclonal anti-SARM1 primary antibody (Chen et al., 2011) followed by Alexa Fluor 488 or 647 anti-mouse secondary antibody (Thermo Fisher Scientific). Fluorescence microscopy images were acquired on a DMi8 upright fluorescence microscope (Leica microsystems) coupled to a monochrome digital camera (Hamamatsu C4742-95). The objective used was HCXPL 20X/0.40 CORR. Numbers of morphologically normal and continuous DsRed labelled neurites and morphologically normal cell bodies were counted in the same field at the indicated timepoints after vacor or vehicle treatment. For each experiment, the average was calculated from three fields per condition; the total number of experiments is indicated in the figure legends. The percentage of healthy neurites and viable neurons remaining relative to the first time point was determined.

### Microfluidic cultures

Dissociated DRG neurons were plated in microfluidic chambers (150 μm barrier, XONA Microfluidics). Cell suspension was pipetted into each side of the upper channel of the microfluidic device. On DIV7, a difference of 100 μl of media between chambers was introduced and drugs were added to the compartment with the lower hydrostatic pressure. To calculate the % of viable neurons, 1 μg/ml propidium iodide (PI) (Thermo Fisher Scientific) was added to the media 15 min before drug addition. Phase contrast and fluorescence microscopy images were acquired on a DMi8 upright fluorescence microscope (Leica microsystems) coupled to a monochrome digital camera (Hamamatsu C4742-95). The objective used was HCXPL 20X/0.40 CORR. For each experiment, the degeneration index was calculated from the average of two distal fields of neurites per condition, whereas the % of viable neurons remaining relative to the first time point was calculated from the average of three fields of cell bodies (staining positive for PI) per condition; the total number of experiments is indicated in the figure legends.

### Intravitreal (IVT) injection into the eye

9-12-week-old male and female C57BL/6J (Charles River, UK) or *Sarm1*^*-/-*^ mice were intravitreally injected with either 2 μL vacor, DMSO (vehicle) or PBS. Injections were performed as previously described (Osborne et al., 2018). ERG responses were recorded simultaneously from both eyes using an Espion E3 system with full-field Ganzfeld sphere (Diagnosys, Cambridge, UK). Animals were dark-adapted overnight, and ERG recordings performed under low level, red light illumination. Peak RGC responses (pSTR) were recorded between 70-110 ms after a light exposure of -4.73 log cd.s.m^-2^, B wave peaks between 80-120 ms, at -1.90 log cd.s.m^-2^, and A wave troughs within the first 20 ms at 1.29 log cd.s.m^-2^. Following culling via exposure to CO_2_, eyes were post-fixed for 24 hr in 4% PFA before retinal flatmounts were prepared as previously described (Osborne et al., 2018). Briefly, retinas were excised from the eye cup, flattened and stained with the RGC specific nuclei marker BRN3A (Santa Cruz, sc-31984, 1:200). Images were captured, blindly, from 8 regions per retina using a 20X objective, and automatically quantified with the ImageJ plugin ‘Simple RGC’ (Cross et al., 2020). The representative images shown were acquired on a confocal microscope (Leica Microsystems) using a 40X objective.

### Determination of NMN, NAD, cADPR, vacor, VMN and VAD tissue levels

Following treatment with vacor or vehicle, DIV7 wild-type and *Sarm1*^−/−^ DRG ganglia were separated from their neurites with a scalpel. Neurite and ganglia (containing short proximal neurite stumps as well as cell bodies) fractions were washed in ice-cold PBS and rapidly frozen in dry ice and stored at −80 °C until processed. Tissues were ground in liquid N_2_ and extracted in HClO_4_ by sonication, followed by neutralisation with K_2_CO_3_. NMN and NAD were subsequently analysed by spectrofluorometric HPLC analysis after derivatisation with acetophenone (Mori et al., 2014). Vacor, VMN and VAD were determined in DRG whole explant cultures by ion pair C18-HPLC chromatography, as previously described (Buonvicino et al., 2018). cADPR levels were determined in DRG whole explant cultures using a cycling assay, as previously described (Graeff and Lee, 2002). A single analysis of vacor, VMN, VAD and cADPR levels was performed in DRG neurite and ganglia fractions independently, which was not further repeated due to the low basal levels of these metabolites and the amount of cellular material needed for this type of analysis. Metabolite levels were normalised to protein levels quantified with the Bio-Rad Protein Assay (Bio-Rad) on formate-resuspended pellets from the aforementioned HClO_4_ extraction.

### VMN and VAD synthesis and purification

VMN and VAD were synthesised as previously reported (Buonvicino et al., 2018) with minor changes. Vacor was either phosphoribosylated *in vitro* by murine NAMPT (mNAMPT) into VMN or phosphoribosylated by mNAMPT and adenylated by murine NMNAT2 (mNMNAT2) into VAD. The scheme of these reactions and the reaction mixtures are shown in Fig. S10A,B. Inorganic yeast pyrophosphatase (PPase), phosphoribosyl 1-pyrophosphate (PRPP) and adenosine triphosphate (ATP) were all from Merck. Recombinant mNAMPT and mNMNAT2 were purified as previously described (Amici et al., 2017; Orsomando et al., 2012). Following incubation, reaction mixtures (A) for VMN and (A+B) for VAD (Fig. S10B) were stopped by rapid cooling on ice and kept refrigerated until injection for purification. Next, the VMN and VAD obtained were purified by FPLC under volatile solvents and lyophilised. Briefly, a preparative IEC chromatography was carried out on AKTA Purifier onto the anion exchanger resin Source 15Q (GE HealthCare, 20 ml volume). The column was equilibrated at room temperature (∼25 °C), at 5 ml/min. After injection of the two mixtures above, a linear gradient elution was applied by mixing the two volatile buffers, as indicated. The eluate was monitored at wavelengths of 260 nm and 340 nm (optimal for vacor nucleotides), and both VMN and VAD peaks were collected (Fig. S10C,D). Next, an additional chromatography was performed by HPLC onto a RP C18 column (Varian, 250 × 4.6 mm, 5μ particles) heated at 50 °C to obtain a pure VAD stock. The equilibration in such case was carried out at 1 ml/min in ammonium formate buffer; this was followed by multiple injections of the previously collected IEC pool of VAD (∼1 ml each), and elution by a linear gradient of increasing acetonitrile in the buffer. The eluate was monitored again at wavelengths 260 nm and 340 nm and the VAD peak was collected (Fig. S10D). After lyophilisation, dry pellets were stored at -80 °C. The resulting VMN and VAD lyophilised powders were 100% pure. Their quantitation after resuspension in H_2_O was performed spectrophotometrically (ɛ_340nm_ of 17.8 mM^-1^cm^-1^).

### Recombinant hSARM1 purification

Recombinant, full-length C-terminal Flag-tagged wild-type and K193R human SARM1 (hSARM1-Flag) were expressed in HEK293T cells and purified by immunoprecipitation. HEK cells at 50-70% confluence were transfected with wild-type and K193R hSARM1-Flag expression construct (pLVX-IRES-ZsGreen vector backbone) using Lipofecatmine 2000 (Thermo Fisher Scientific). To boost wild-type and K193R hSARM1-Flag expression, media were supplemented with 2 mM nicotinamide riboside (prepared from Tru Niagen^®^ capsules by dissolving the contents and passing through a 0.22 µm filter) at the time of transfection. After 24 hr, cells were collected and washed in cold PBS and lysed for 10 min with trituration by pipetting and repeated vortexing in ice-cold KHM buffer (110 mM potassium acetate, 20 mM HEPES pH 7.4, 2 mM MgCl_2_, 0.1 mM digitonin) with cOmplete™, Mini, EDTA-free protease inhibitor cocktail (Roche). After centrifugation for 5 min at 3000 rpm in a chilled microfuge, the supernatant was collected and diluted to 1 µg/µl in KHM buffer after determination of its protein concentration by the Pierce BCA assay (Thermo Fisher Scientific). For immunoprecipitation, 1 ml of extract was incubated overnight at 4°C with rotation with 20 µg/ml anti-FLAG M2 antibody (Merck, F3165) and 50 µl Pierce magnetic protein A/G beads (Thermo Fisher Scientific) pre-washed with KHM buffer. Beads were collected on a magnetic rack and washed 3x with 500 µl KHM buffer and 1x with PBS (with protease inhibitors) and then resuspended in PBS containing 1 mg/ml BSA. hSARM1-Flag concentration in the bead suspension was determined relative to an hSARM1 standard (purified from insect cells) by immunoblotting using a rabbit polyclonal antibody raised against human SAM-TIR.

### hSARM1 NADase activity

Rates of NAD consumption by recombinant wild-type and K193R hSARM1 were measured by HPLC under a discontinuous assay that was set as follows. Typically, mixtures of 0.02-0.2 ml contained 2-20 μg/ml of hSARM1-Flag (on-beads) in buffer HEPES/NaOH 50 mM, pH 7.5, and 250 μM NAD. NMN, VMN and VAD were added at the concentrations indicated in the figures. Reactions were initiated by adding NAD, incubated at 25 °C in a water-bath, stopped at appropriate times by HClO_4_ treatment, neutralised with K_2_CO_3_, and subsequently analysed by ion-pair reverse-phase (RP) HPLC (NAM, NMN, NAD, ADPR, cADPR) (Mori et al., 2014) or under optimised conditions for VMN and VAD detection (Buonvicino et al., 2018). The products formed (NAM, ADPR and cADPR) were quantified from the area of separated peaks. Rates were calculated under NAD consumption ≤ 20% from the linearly accumulating products ADPR and cADPR (Fig. S6B). One unit (U) of activity represents the enzyme amount that forms 1 μmol/min of the products above under these assay conditions. Data from three independent experiments were averaged to calculate hSARM1 activity fold activation (Fig. 4A) and then studied by fitting through the Excel software package to the rate equation below

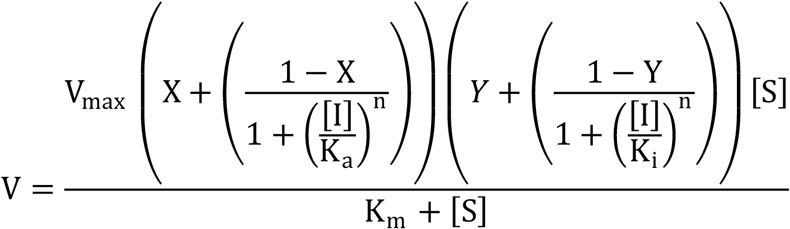

that represents a Michaelis-Menten equation adapted to be valid for readily reversible effectors, *i*.*e*. including activators and inhibitors binding via distinct sites to one protein target. Maximum velocity (*V*_max_) represents the rate in the absence of the effector (basal activity) corrected by the two parameters between parentheses, the first for activation and the second for inhibition, taking into account opposite and independent contributions exerted by the single but dual effector I. *X* is the “relative activity fold activation” factor at saturating concentrations of I, *Y* is the residual activity fraction at saturating concentration of I, *K*_*a*_ is the affinity constant of I for the activation site, *K*_*i*_ is the affinity constant of I for the inhibition site, and *n* is the Hill coefficient indicating cooperativity (if ≠ 1).

### Expression and purification of recombinant dSARM1^ARM^

Codon-optimised cDNA of dSARM1^ARM^ (residues 315-678) was cloned into the pMCSG7 expression vector at the SspI site, using ligation-independent cloning (Aslanidis and de Jong, 1990). Plasmids encoding various dSARM1^ARM^ mutants were generated using the Q5^®^ Site-Direct Mutagenesis Kit (New England BioLabs). Plasmids were transformed into BL21 (DE3) cells. Large-scale expression of dSARM1^ARM^ and its mutants was performed using the auto-induction method (Studier, 2005). In brief, 2 ml of overnight culture of the transformed BL21 cells was inoculated into 1 l of auto-induction media with 100 μg/ml ampicillin, and grew at 37 °C, 225 rpm to reach OD_600_ of 0.8-1.0. Temperature was decreased to 20 °C for protein expression overnight. Cells were harvested by centrifugation at 6000×g for 20 min at 4 °C, and resuspended into lysis buffer (50 mM HEPES pH 8.0, 500 mM NaCl and 30 mM imidazole). PMSF (phenylmethanesulfony fluoride) was added to the cell resuspension at 1 mM concentration. Cell resuspension from 1 l of culture was sonicated at the amplitude of 40% for 120 s (10 s on and 20 s off). Lysed cells were centrifuged at 15300×g for 40 min at 4 °C. The supernatant was loaded onto the HisTrap HP 5 ml Ni column, followed by washing using 10 CV of lysis buffer. The bound protein was eluted using elution buffer (50 mM HEPES pH 8.0, 500 mM NaCl and 300 mM imidazole), and incubated with TEV (tobacco etch virus) protease to remove the N-terminal 6×histidine tag in the SnakeSkin Dialysis Tubing, 3.5K MWCO (Thermo Fisher Scientific), dialysed against the buffer containing 20 mM HEPES (pH 8.0), 300 mM NaCl and 1 mM DTT at 4 °C overnight. The cleaved protein was reloaded onto the Ni column and flow-through was collected, concentrated to 10 ml and injected onto the Superdex 75 HiLoad 26/600 column (GE Healthcare) equilibrated with gel-filtration buffer (10 mM HEPES pH 8.0 and 150 mM NaCl). The pure target protein, as confirmed by SDS-PAGE analysis and mass spectrometry, was concentrated to 10 mg/ml and stored at -80 °C.

### Protein crystallisation, diffraction data collection and structural determination

dSARM1^ARM^ (10 mg/ml) and VMN were incubated at 1: 10 molar ratio at 4 °C overnight. Hanging drops, containing 2 μl of protein complex and 2 μl of well solution (20 % PEG 3350 and 0.2 M magnesium acetate tetrahydrate, pH 7.9), were equilibrated against 500 μl of well solution at 20 °C. Crystals with irregular shapes were observed after 3-5 days. *In situ* partial proteolysis happened during the process of crystallisation, as the analysis of the crystals via SDS-PAGE analysis and mass spectrometry indicated that the crystallised protein only spanned residues 370-678, rather than residues 315-678. Crystals cryo-protected in paratone-N diffracted to 1.7-2.3 Å resolution at the Australian Synchrotron MX2 beamline. X-ray diffraction data was collected at the wavelength of 0.95372 Å. Raw data was processed using XDS (Kabsch, 2010); data reduction, scaling and reflection selection for R_free_ calculation were performed using Aimless within CCP4 suite (Evans and Murshudov, 2013). Phases were calculated via molecular replacement using the NMN-bound dSARM1^ARM^ (PDB: 7LCZ) (Figley et al., 2021; Gu et al., 2021) as a search model in the program Phaser within Phenix (McCoy et al., 2007). The model was refined using COOT (Emsley and Cowtan, 2004) and Phenix (Afonine et al., 2012), and analysed using PyMol (Schrodinger), PDBsum (Laskowski et al., 2018) and PISA (Krissinel and Henrick, 2007). The crystals contain two molecules of VMN-bound dSARM1^ARM^ (RMSD = 0.3 Å over 296 Cα atoms) in the asymmetric unit.

### Isothermal titration calorimetry (ITC)

ITC experiments were performed in duplicate on Nano ITC (TA Instruments). All proteins and compounds were dissolved in a buffer containing 10 mM HEPES (pH 8.0) and 150 mM NaCl. The baseline was equilibrated for 600 s before the first injection. VMN (0.6 mM) was titrated as 20-25 injections of 1.96 μL every 200 s, into 45 μM protein. The heat change was recorded by injection over time and the binding isotherms were generated as a function of molar ratio of the protein solution. The dissociation constants (K_d_) were obtained after fitting the integrated and normalised data to a single-site binding model using NanoAnalyze (TA Instruments).

### NMN deamidase activity

Recombinant *E. coli* NMN deamidase was obtained as described previously (Zamporlini et al., 2014). Activity was measured in buffer HEPES/NaOH 50 mM, pH 7.5, in the presence of 4 milliU/ml enzyme, 0.5 mg/ml BSA, and 250 μM NMN or VMN. Vacor, VMN and VAD, all at the concentration of 250 μM, were also assayed in presence of the substrate NMN. Reactions were incubated at 37 °C, then stopped and analysed by HPLC using the two methods described above (Buonvicino et al., 2018; Mori et al., 2014). The NMN deamidase rates were calculated after separation and quantification of the NaMN product formed from NMN, and finally reported as relative percentages of controls in the presence of NMN alone.

### Western blot

Following treatment with vacor, DRG ganglia were separated from their neurites with a scalpel. Neurites and ganglia were collected, washed in ice-cold PBS containing protease inhibitors (Roche), and lysed directly in 15 µl 2x Laemmli buffer containing 10% 2-mercaptoethanol (Merck). Samples were loaded on a 4-to-20% SDS polyacrylamide gel (Bio-Rad). Membranes were blocked for 3 hr in 5% milk in TBS (50 mM Trizma base and 150 mM NaCl, PH 8.3, both Merck) plus 0.05% Tween-20 (Merck) (TBST), incubated overnight with primary antibody in 5% milk in TBST at 4°C and subsequently washed in TBST and incubated for 1 hr at room temperature with HRP-linked secondary antibody (Bio-Rad) in 5% milk in TBST. Membranes were washed, treated with SuperSignal™ West Dura Extended Duration Substrate (Thermo Fisher Scientific) and imaged with Uvitec Alliance imaging system. The following primary antibodies were used: mouse monoclonal anti-SARM1 (Chen et al., 2011) (1:5000) and mouse anti-β-actin (Merck, A5316, 1:2000) as a loading control. Quantification of band intensity was determined by densitometry using ImageJ.

### Statistical analysis

Statistical testing of data was performed using Prism (GraphPad Software, La Jolla, USA). The appropriate tests used and the n numbers of each individual experiment are described in the figure legends. A p-value < 0.05 was considered significant.

## Data and materials availability

The data that support the findings of this study are available from the corresponding author upon reasonable request. VMN-bound dSARM1^ARM^ crystal structure has been deposited in the Protein Data Bank (PDB: 7M6K).

## Acknowledgments

We thank members of the Coleman lab, Kobe lab, Prof James Fawcett and Prof Nadia Raffaelli for useful discussions. We also thank Dr Lucia Silvestrini for help in obtaining the Neurospora crassa NADase for the cADPR detection assay. We acknowledge the use of the University of Queensland Remote Operation Crystallization and X-ray (UQROCX) Facility at the Centre for Microscopy and Microanalysis and the Australian Synchrotron MX beamlines, and the staff for support. This work was funded by a Sir Henry Wellcome postdoctoral fellowship from the Wellcome Trust [grant number 210904/Z/18/Z], a Wellcome Trust Clinical Research Career Development Fellowship [grant number: 206634]; BBSRC/AstraZeneca Industrial Partnership Award BB/S009582/1; Grants RSA 2016-18 and 2017-19 from UNIVPM (Ancona, Italy); Australian National Health and Medical Research Council (NHMRC 1160570); Australian Research Council (ARC; Laureate Fellowship FL180100109) to B.K. and National Eye Research Centre (SAC 041). This research was funded in whole, or in part, by the Wellcome Trust [Grant number 210904/Z/18/Z and 206634]. For the purpose of open access, the author has applied a CC BY public copyright licence to any Author Accepted Manuscript version arising from this submission.

## Author contributions

A.L., G.O. and M.P.C. conceived the study. A.L. designed and performed all experiments in neurons, with help from E.M.; C.A. and G.O. designed and performed nucleotide measurements and biochemical assays, with help from A.A.; A.O. and B.N. performed *in vivo* experiments. W.G., Z.L. and B.K. determined and analysed the crystal structure of VMN bound to dSARM1 ARM domain. T.V. and L. H-T. performed ITC experiments. J.G. contributed to the interpretation of the results and developed the hSARM1 purification protocol, with help from L.M.D.; P.A-F. and Q.W. contributed to the interpretation of the results. A.L. and M.P.C. wrote the manuscript, with input from J.G., A.O., W.G., B.K. and G.O. All authors read and approved the manuscript.

