## Supplemental data for "Potent activation of SARM1 by NMN analogue VMN underlies vacor neurotoxicity"

Fig. S1

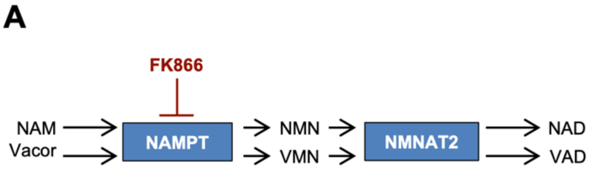


B

4 hours after Vacor addition

| | pmol/µg protein | SEM +/- |
|---|---|---|
| **Vacor** | 2.10 | 0.281 |
| **VMN** | 0.17 | 0.039 |
| **VAD** | 0.14 | 0.017 |

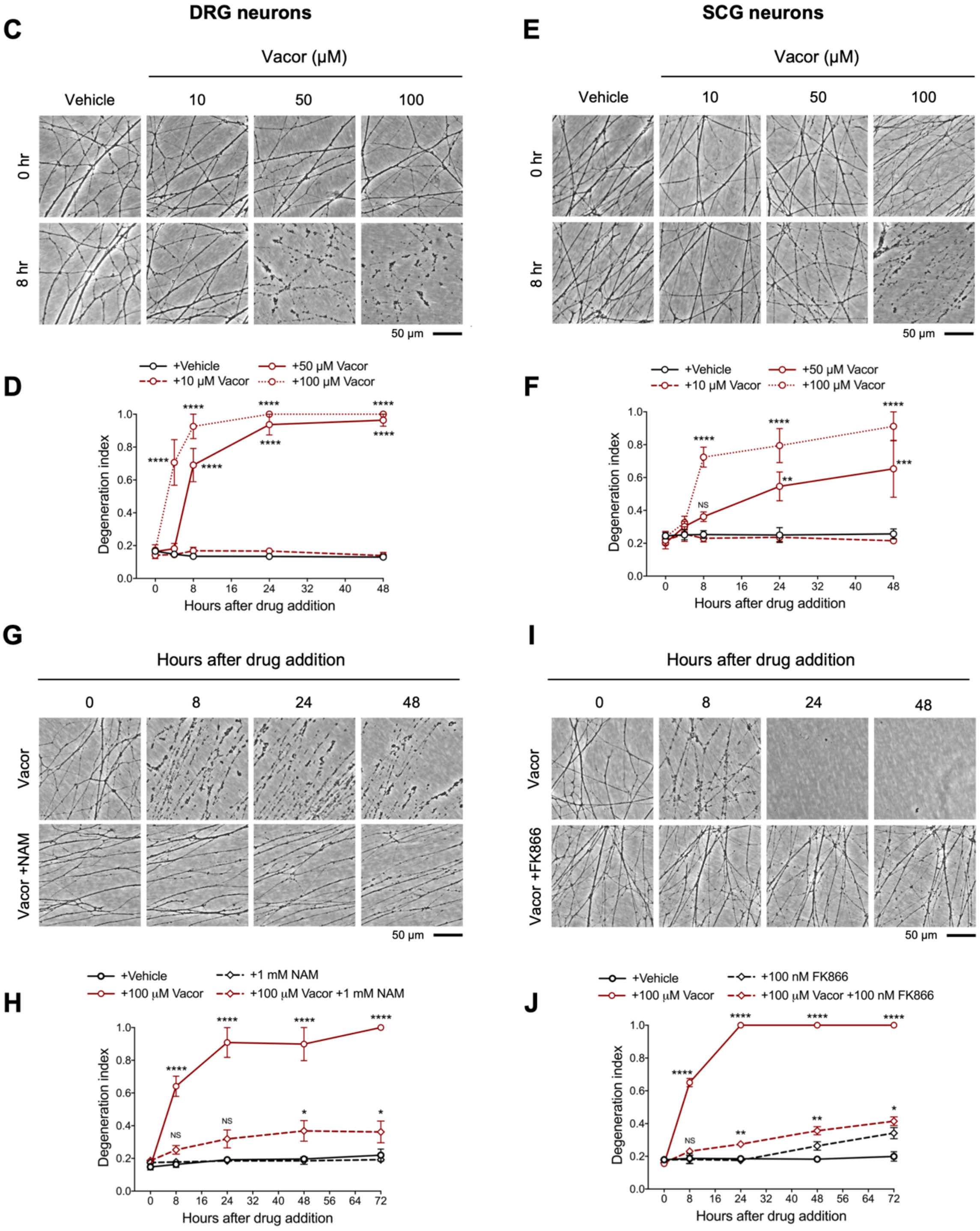

**Fig. S1. Related to Fig. 1. Vacor causes neurite degeneration in primary mouse neurons.**

**(A)** Schematic representation of vacor conversion into VMN and VAD by NAMPT and NMNAT2, respectively. VAD has been reported to inhibit NMNAT2 (Buonvicino et al., 2018). Vacor competes with NAM for NAMPT. High doses of NAM or inhibition of NAMPT with FK866 prevent vacor conversion into downstream metabolites (NAM, nicotinamide; NaMN, nicotinic acid mononucleotide; NMN, nicotinamide mononucleotide; NAD, nicotinamide adenine dinucleotide; NAMPT, nicotinamide phosphoribosyltransferase; NMNAT2, nicotinamide mononucleotide adenylyltransferase 2; VMN, vacor mononucleotide; VAD, vacor adenine dinucleotide). **(B)** Vacor, VMN and VAD levels in wild-type DRG whole explant cultures (neurites and cell bodies) 4 hr after 50 μM vacor treatment (mean ± SEM; n = 3). **(C)** Representative images of neurites from wild-type DRG explant cultures treated with 10, 50, 100 μM vacor or vehicle. **(D)** Quantification of the degeneration index in experiments described in (C) (mean ± SEM; n = 3; repeated measures two-way ANOVA followed by Tukey's multiple comparison test; ****, $p < 0.0001$; statistical significance shown relative to +Vehicle). **(E)** Representative images of neurites from wild-type SCG explant cultures treated with 10, 50, 100 μM vacor or vehicle. **(F)** Quantification of the degeneration index in experiments described in (E) (mean ± SEM; n = 3; repeated measures two-way ANOVA followed by Tukey's multiple comparison test; ****, $p < 0.0001$; ***, $p < 0.001$; **, $p < 0.01$; NS, not-significant; statistical significance shown relative to +Vehicle). **(G)** Representative images of neurites from wild-type SCG explant cultures treated with 100 μM vacor or 100 μM vacor + 1 mM NAM. **(H)** Quantification of the degeneration index in experiments described in (G) (mean ± SEM; n = 3; repeated measures two-way ANOVA followed by Tukey's multiple comparison test; ****, $p < 0.0001$; *, $p < 0.05$; NS, not-significant; statistical comparisons shown are: +100 μM Vacor vs +100 μM Vacor +1 mM NAM and +1 mM NAM vs +100 μM Vacor +1 mM NAM). **(I)** Representative images of neurites from wild-type SCG explant cultures treated with 100 μM vacor, 100 nM FK866 or vehicle. **(J)** Quantification of the degeneration index in experiments described in (I) (mean ± SEM; n = 3; repeated measures two-way ANOVA followed by Tukey's multiple comparison test; ****, $p < 0.0001$; **, $p < 0.01$; *, $p < 0.05$; NS, not-significant; statistical comparisons shown are: +100 μM Vacor vs +100 μM Vacor +100 nM FK866 and +100 nM FK866 vs +100 μM Vacor +100 nM FK866).

Fig. S2

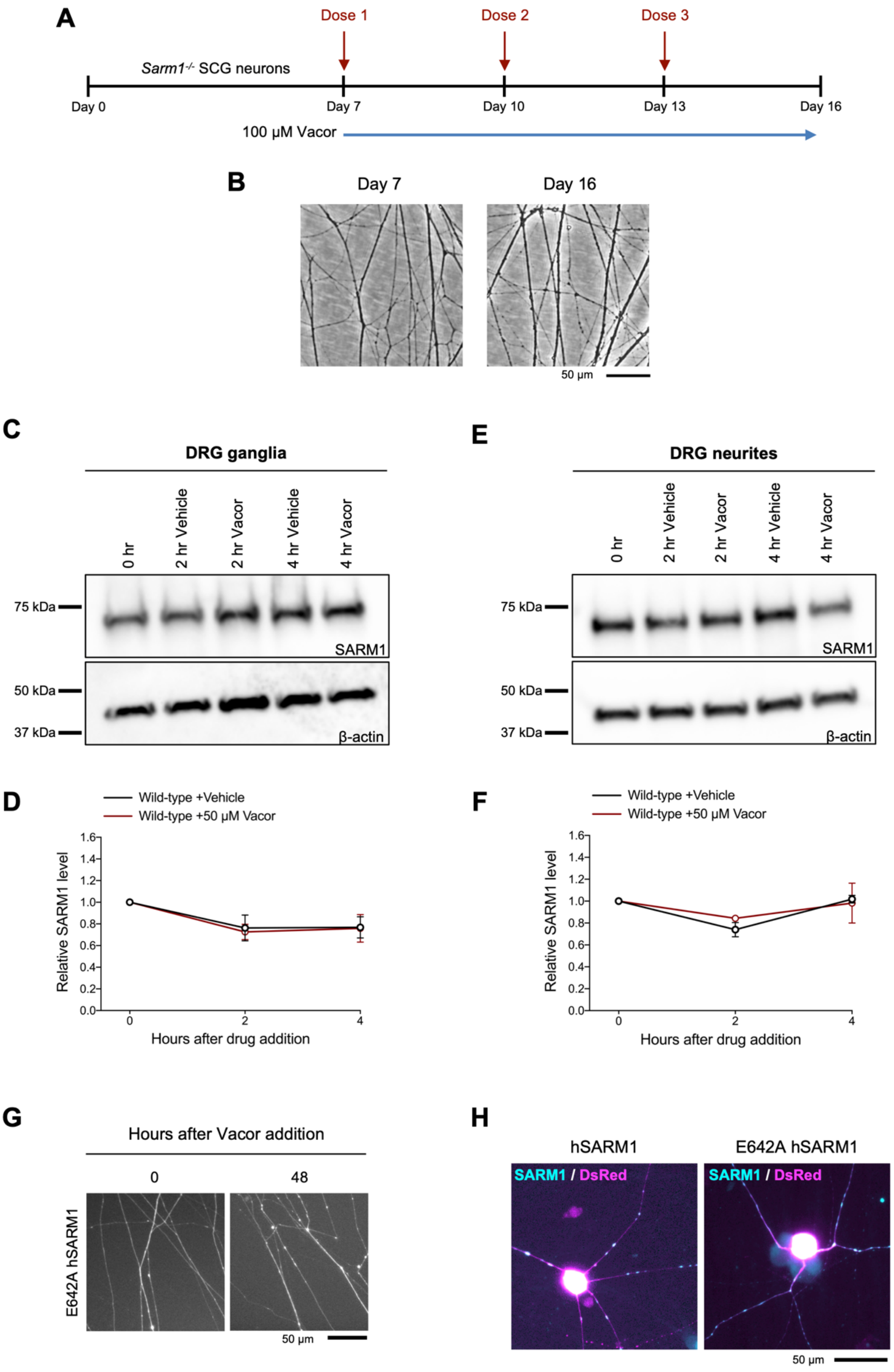
A
Dose 1
Dose 2
Dose 3
Sarm1-/- SCG neurons
Day 0
Day 7
Day 10
Day 13
Day 16
100 µM Vacor
B
Day 7
Day 16
50 µm
C
DRG ganglia
0 hr
2 hr Vehicle
2 hr Vacor
4 hr Vehicle
4 hr Vacor
75 kDa
SARM1
50 kDa
37 kDa
β-actin
E
DRG neurites
0 hr
2 hr Vehicle
2 hr Vacor
4 hr Vehicle
4 hr Vacor
75 kDa
SARM1
50 kDa
37 kDa
β-actin
D
Wild-type +Vehicle
Wild-type +50 µM Vacor
Relative SARM1 level
Hours after drug addition
F
Wild-type +Vehicle
Wild-type +50 µM Vacor
Relative SARM1 level
Hours after drug addition
G
Hours after Vacor addition
0
48
E642A hSARM1
50 µm
H
hSARM1
E642A hSARM1
SARM1 / DsRed
SARM1 / DsRed
50 µm

**Fig. S2. Related to Fig. 1. Long-term survival of *Sarm1$^{-/-}$* SCG neurites following multiple vacor doses.**

**(A)** Schematic representation of the experimental design. *Sarm1$^{-/-}$* SCG explant cultures were treated with 100 μM vacor at DIV7. Multiple doses of vacor were administered by replacing the media every 3 days with fresh media containing vacor. **(B)** Representative images of neurites from *Sarm1$^{-/-}$* SCG explant cultures showing no degeneration at 16 days, after treatment with three doses of 100 μM vacor. **(C)** Representative immunoblots of the ganglia fraction from wild-type DRG explant cultures at the indicated time points after 50 μM vacor or vehicle probed for SARM1 and β-actin (loading control). **(D)** Quantification of normalised SARM1 level (to β-actin) is shown, with data presented relative to 0 hr (mean ± SEM; n = 4; two-way ANOVA followed by Tukey's multiple comparison test). **(E)** Representative immunoblots of the neurite fraction from wild-type DRG explant cultures at the indicated time points after 50 μM vacor or vehicle probed for SARM1 and β-actin (loading control). **(F)** Quantification of normalised SARM1 level (to β-actin) is shown, with data presented relative to 0 hr (mean ± SEM; n = 4; two-way ANOVA followed by Tukey's multiple comparison test). **(G)** Representative images showing long-term survival of neurites from *Sarm1$^{-/-}$* SCG dissociated neurons co-injected with plasmids encoding E642A hSARM1 and DsRed (to label neurites) and treated with 100 μM vacor. **(H)** Representative images showing expression of both wild-type and E642A hSARM1 for microinjection experiments in (Fig. 1G-I). The absence of vacor-induced toxicity in neurons expressing E642A hSARM1 is therefore due to loss of enzymatic activity, rather than the lack of protein expression.

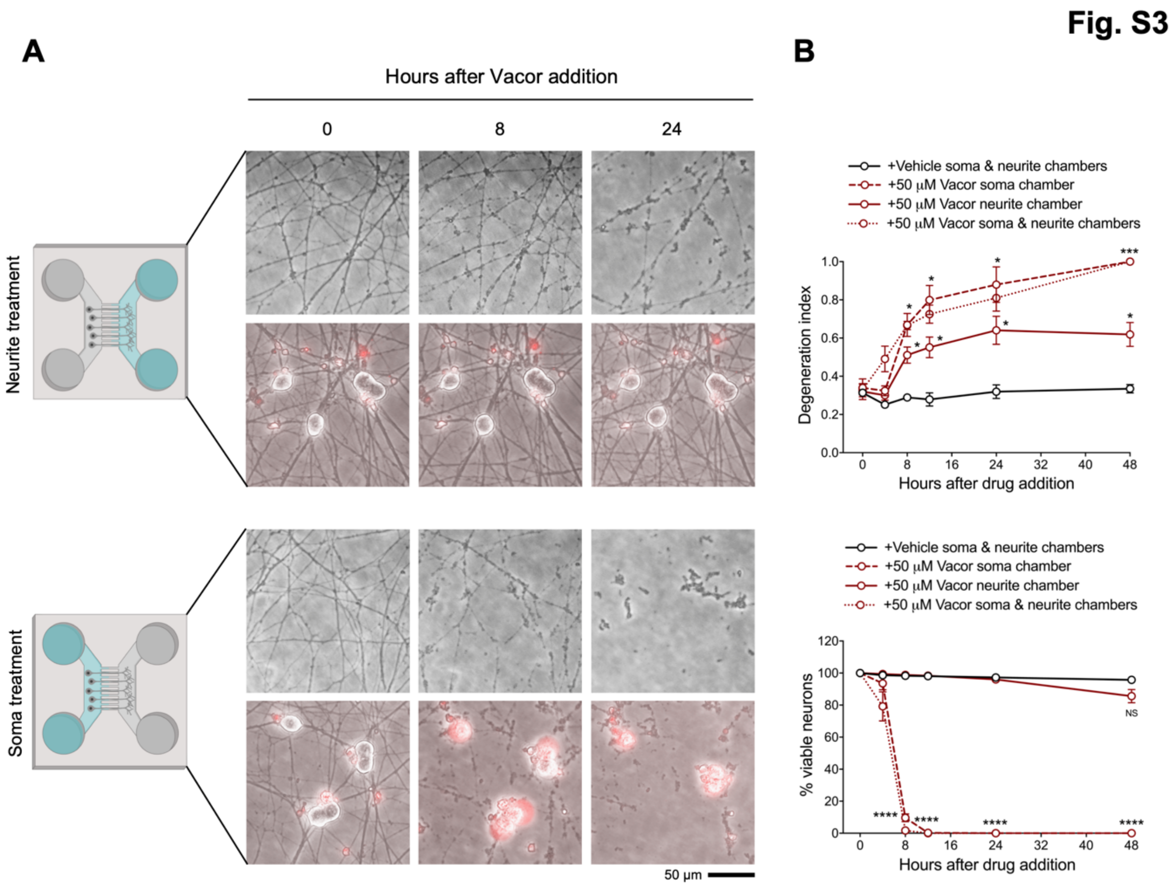


**Fig. S3. Related to Fig. 1. Local death of neurites and cell bodies caused by vacor.**

**(A)** Representative images of neurites and cell bodies from wild-type DRG dissociated neurons cultured in microfluidic chambers and treated with 50 µM vacor or vehicle ('Created with BioRender'). **(B)** Quantification of the degeneration index and viable neurons (shown as a percentage relative to 0 hr) in experiments described in (A) (mean ± SEM; n = 4; repeated measures two-way ANOVA followed by Tukey's multiple comparison; ***, $p < 0.001$; *, $p < 0.05$; statistical significance shown relative to +Vehicle soma & neurite chambers).

Fig. S4

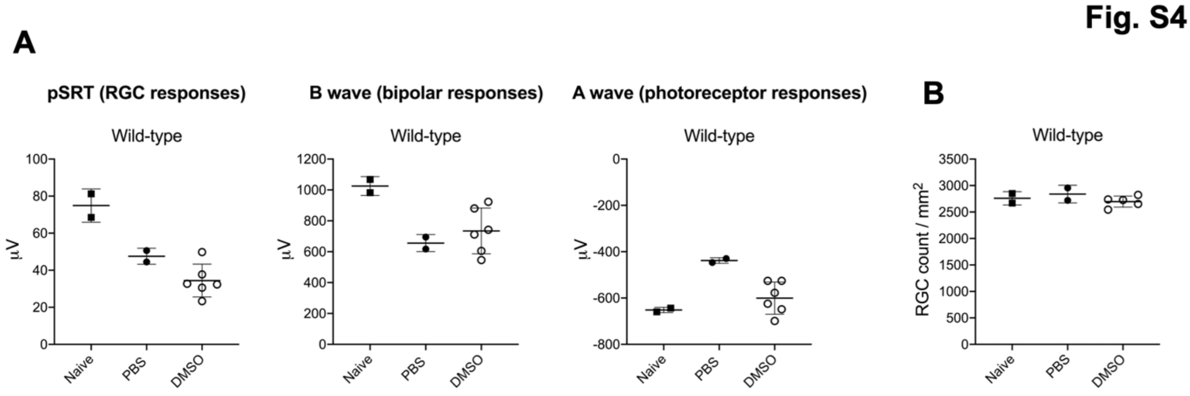


**Fig. S4. Related to Fig. 2. DMSO has no adverse effect on retinal cell survival or function compared to PBS-injected eyes.**

**(A)** Quantification of the ERG responses (pSTR, B wave and A wave) from wild-type mice untreated (naive-no injection) or injected with PBS or DMSO (mean ± SD; n = 2-6). **(B)** Quantification of RGC numbers from wild-type mice untreated (naive-no injection) or injected with PBS or DMSO (mean ± SD; n = 2-5).

Fig. S5

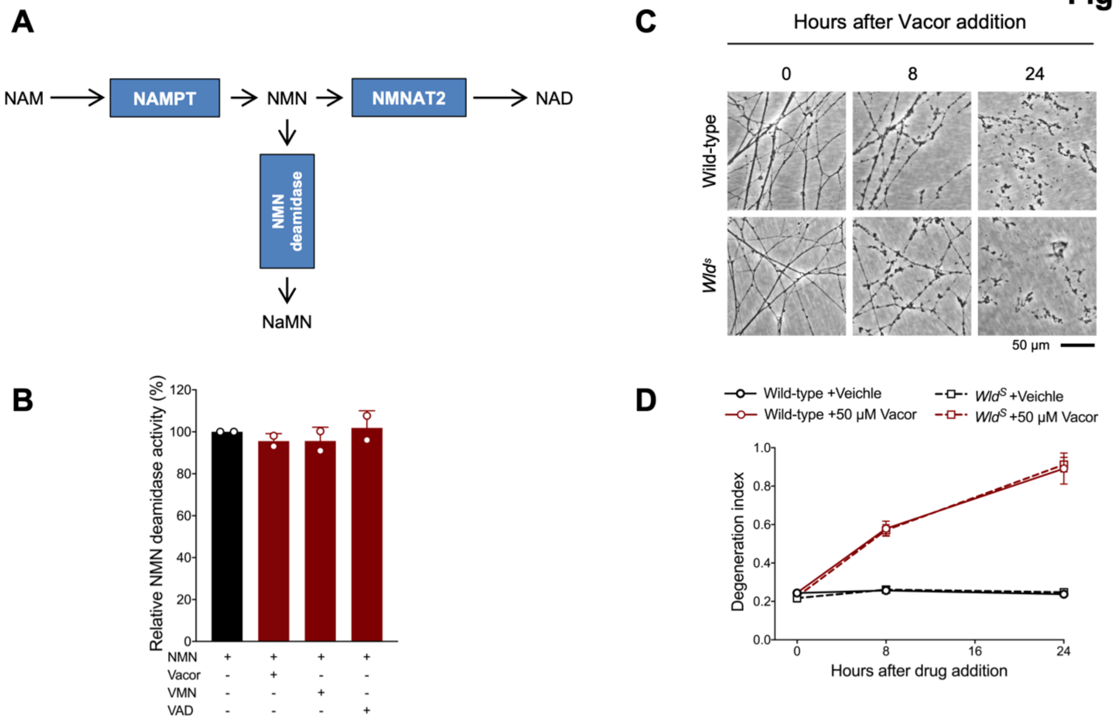


**Fig. S5. Related to Fig. 3. Effect of vacor, VMN and VAD on recombinant NMN deamidase activity and lack of protection after vacor treatment in *Wld*$^{S}$ neurons.**

**(A)** Schematic representation of NAD biosynthetic pathway from NAM and the side reaction catalysed by bacterial NMN deamidase, which prevents accumulation of the NMN intermediate. **(B)** Relative activity (%) (NMN conversion into NaMN) of purified, recombinant NMN deamidase in the presence of vacor, VMN and VAD (all 250 µM) (mean ± SD; n = 2). **(C)** Representative images of neurites from wild-type and *Wld*$^{S}$ DRG explant cultures treated with 50 µM vacor. **(D)** Quantification of the degeneration index in experiments described in (C) (mean ± SEM; n = 4; repeated measures three-way ANOVA followed by Tukey's multiple comparison test).

Fig. S6

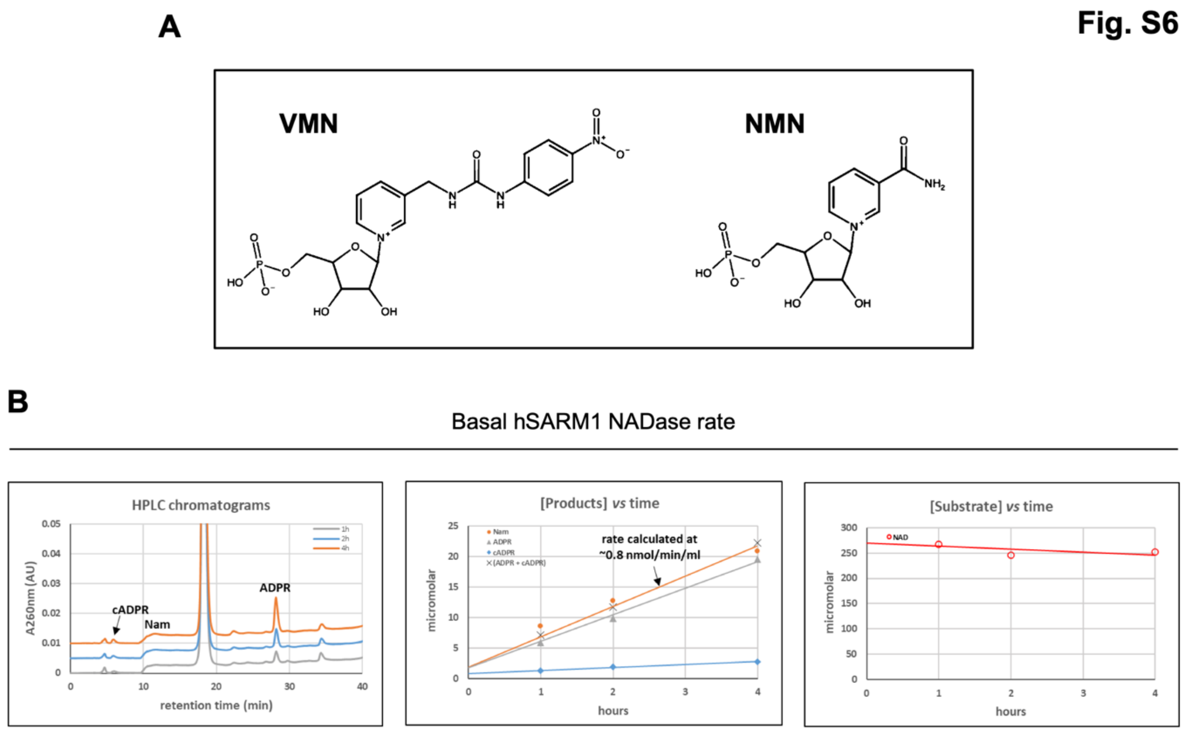


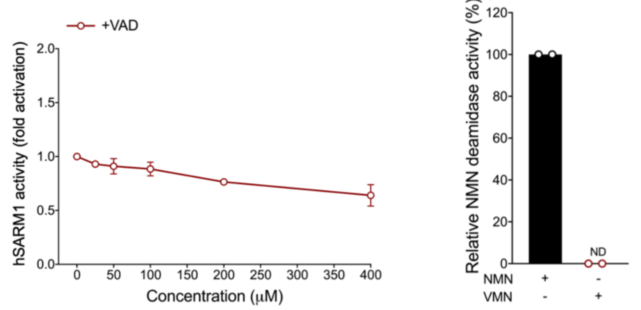


**E**

4 hours after Vacor addition

| | DRG ganglia | DRG neurites |
|---|---|---|
| | pmol/μg protein | pmol/μg protein |
| **Vacor** | 2.25 | 1.09 |
| **VMN** | 0.17 | 0.02 |
| **VAD** | 0.16 | 0.03 |

**Fig. S6. Related to Fig. 4. Representative basal NADase rate of recombinant hSARM1.**

**(A)** Chemical structures of the two pyridine 5' mononucleotides VMN and NMN. **(B)** Representative chromatogram and graphs showing basal NADase activity of purified, recombinant hSARM1 (40 μg/ml). The formed products after separation (left panel) are quantified by peak-area integration and evaluated for linearity (middle panel). The NADase rate is calculated from accumulating products ADPR and cADPR (middle panel). In every experiment, the sum of ADPR and cADPR fully matched the amount of NAM formed (middle panel), as well as the amount of NAD consumed (right panel). **(C)** Fold change of NADase activity of purified, recombinant hSARM1 in the presence of VAD (mean ± SD; n = 2). Rates are relative to control measured with 250 μM NAD alone. VAD, once added to the reaction mixture, was not consumed during incubation.

**(D)** Relative activity (%) of purified, recombinant NMN deamidase with VMN as a substrate. NMN or VMN were assayed at 250 µM (mean ± SD; n = 2). **(E)** A single analysis of vacor, VMN and VAD levels in wild-type DRG ganglia and neurite fractions 4 hr after 50 µM vacor treatment (n = 1).

Fig. S7

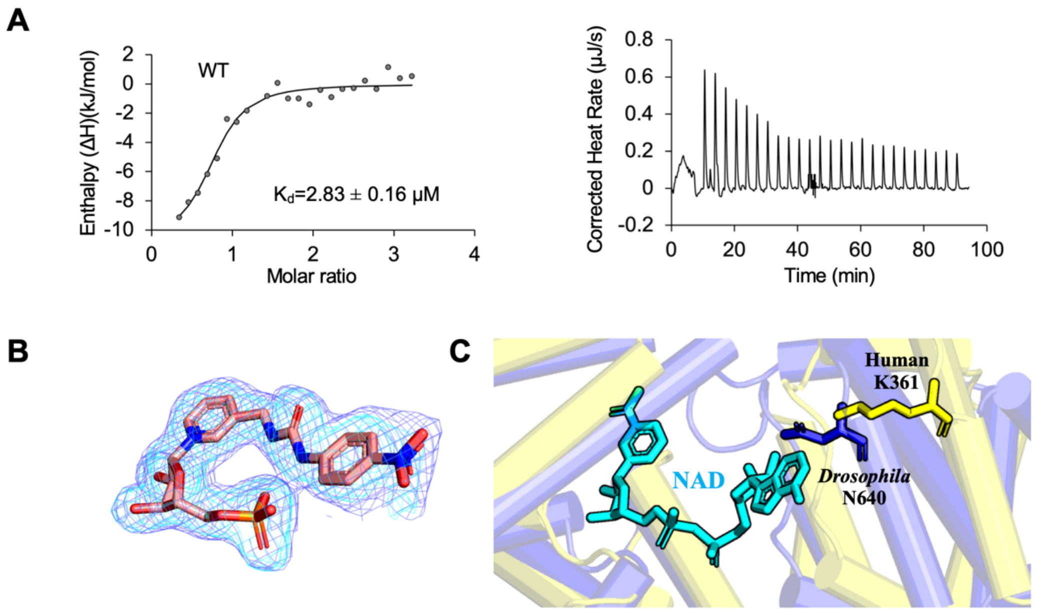


**Fig. S7. Related to Fig. 4. Analysis of dSARM1$^{ARM}$ : VMN interaction.**

**(A)** Integrated (left) and raw (right) ITC data for the titration of 0.6 mM VMN with 45 μM dSARM1$^{ARM}$. **(B)** Standard omit (cyan) and Polder (slate) mFo-DFc maps near the VMN molecule in dSARM1$^{ARM}$ crystals (chain A). **(C)** The adenine group of NAD clashes with *Drosophila* N640 (human K361) in the VMN-bound structure.

**Fig. S8**

**A**

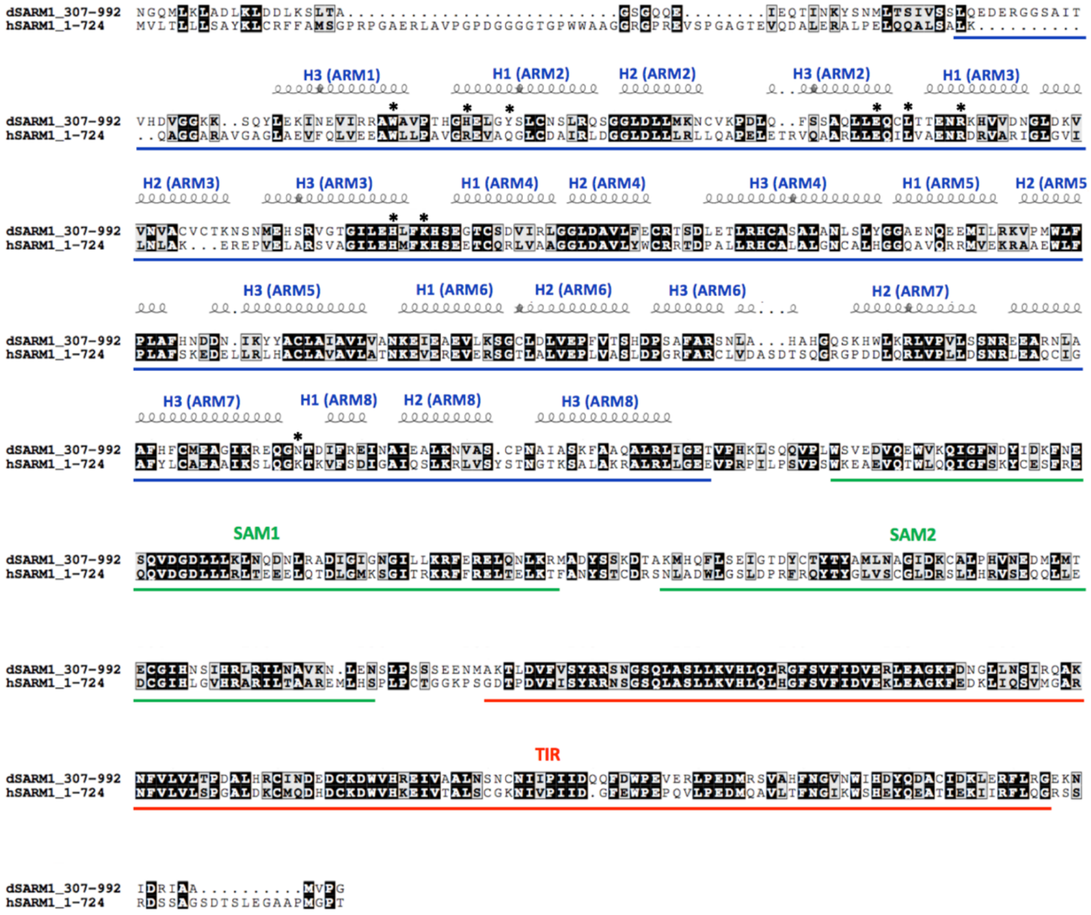


**B**

| ***Drosophila*** (Human) | VMN moieties | Interaction |
|---|---|---|
| **W385** (W103) | pyridine | pi-stacking |
| **H392** (R110) | nitrobenzene | pi-stacking |
| | urea | hydrogen bond |
| **Y396** (Q114) | phosphate | hydrogen bond |
| **E429** (E149) | ribose | hydrogen bond |
| **L432** (L152) | ribose | hydrogen bond |
| **R437** (R157) | phosphate | hydrogen bond |
| **H473** (H190) | ribose | hydrogen bond |
| **K476** (K193) | phosphate | hydrogen bond |
| **N640** (K361) | nitro | hydrogen bond |

**Fig. S8. Related to Fig. 4. Sequence alignment of SARM1 orthologs.**

**(A)** Sequence alignment of *Drosophila* and human SARM1. The alignment was performed and analysed using T-coffee Multiple Sequence Alignment Server (Expresso) (Notredame et al., 2000) and ESPript, respectively. Conserved residues are highlighted in black boxes. Residues important for VMN interaction are indicated by stars. **(B)** Detailed interaction between VMN and the residues indicted by stars in (A).

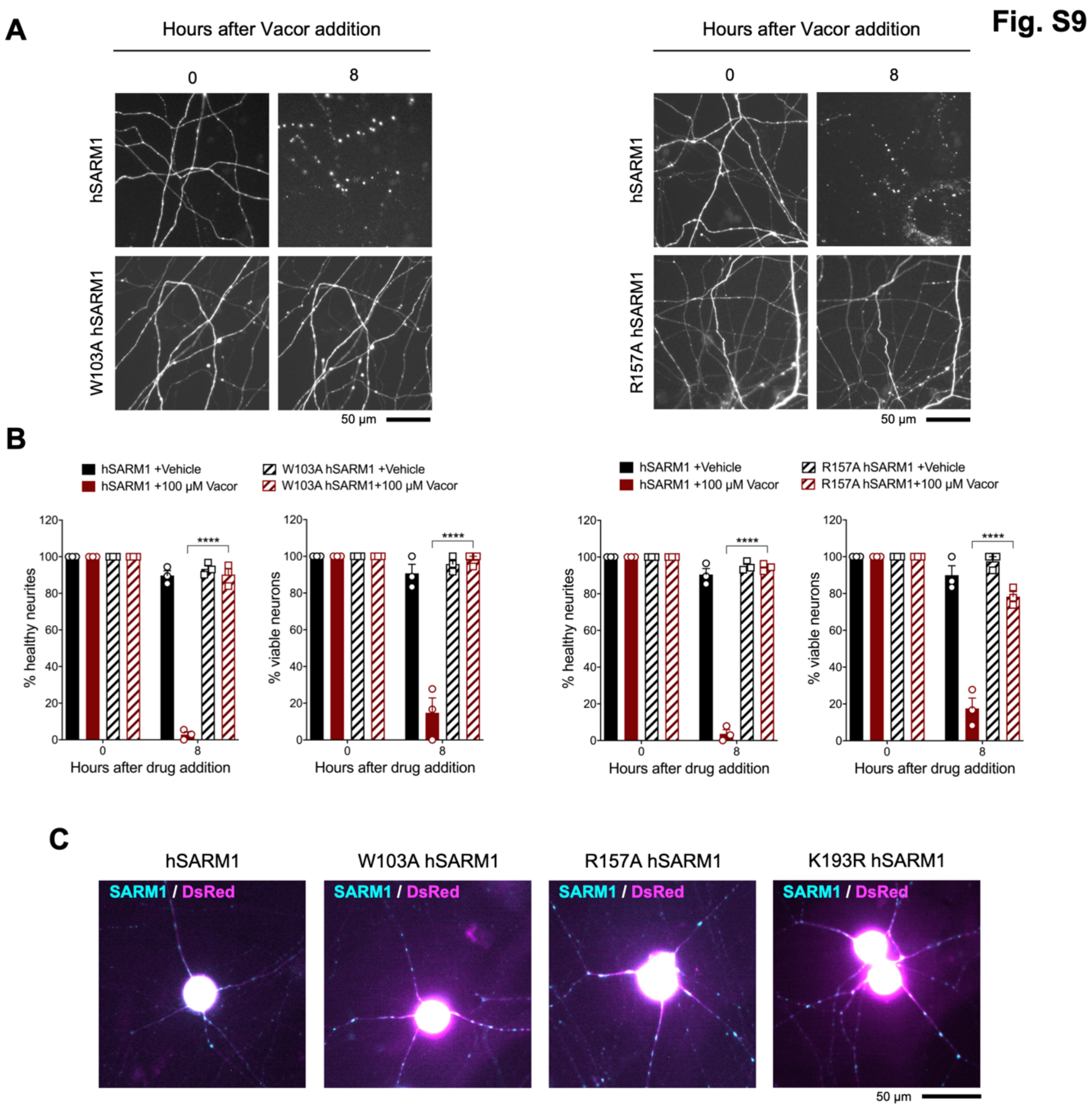


**Fig. S9. Related to Fig. 4. Mutations in the VMN binding pocket of hSARM1 ARM domain prevent vacor toxicity.**

**(A)** Representative images of neurites from *Sarm1*$^{-/-}$ SCG dissociated neurons co-injected with plasmids encoding wild-type, W103A or R157A hSARM1 and DsRed (to label neurites) and treated with 100 μM vacor. **(B)** Quantification of healthy neurites and viable neurons in experiments in (A) is shown as a percentage relative to 0 hr (time of drug addition) (mean ± SEM; n = 3; repeated measures three-way ANOVA followed by Tukey's multiple comparison test; ****, $p < 0.0001$). **(C)** Representative images showing expression of wild-type, W103A, R157A and K193R hSARM1 for microinjection experiments in Fig. 4G,H and Fig. S9A,B.

Fig. S10

A

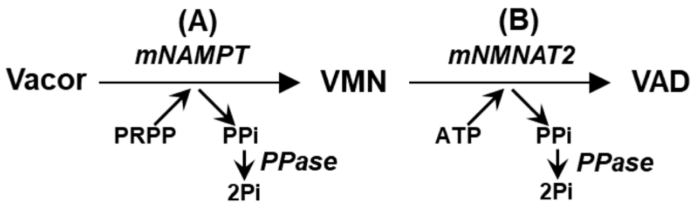


B

| | Mix (A) for VMN | | Mix (A+B) for VAD | |
|---|---|---|---|---|
| | ml | final [ ] | ml | final [ ] |
| $H_2O$ | 12.8 | - | 12.2 | |
| HEPES/NaOH 1 M, pH 7.5 | 0.75 | 50 mM | 0.75 | 50 mM |
| $MgCl_2$ 1 M | 0.30 | 20 mM | 0.30 | 20 mM |
| ATP 100 mM | 0.15 | 1 mM | 0.30 | 2 mM |
| PRPP 50 mM | 0.15 | 0.5 mM | 0.15 | 0.5 mM |
| Vacor 9 mM | 0.50 | 0.3 mM | 0.50 | 0.3 mM |
| mNAMPT wild type ~0.25 U/ml | 0.35 (*) | ~6 mU/ml | 0.35 (*) | ~0.006 U/ml |
| mNMNAT2 wild type ~2.5 U/ml | - | - | 0.44 (*) | ~0.075 U/ml |
| Yeast PPase Sigma >2 KU/ml | - | - | 0.01 | >1.33 U/ml |
| Final mix | 15 ml | | 15 ml | |
| Incubation settings | 2-3 hr at 37 °C | | Overnight (~19 hr) at 30 °C | |

(*) Reagent(s) added to start the reactions.

C

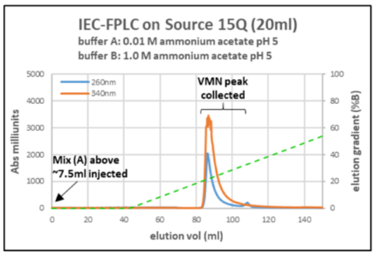


D

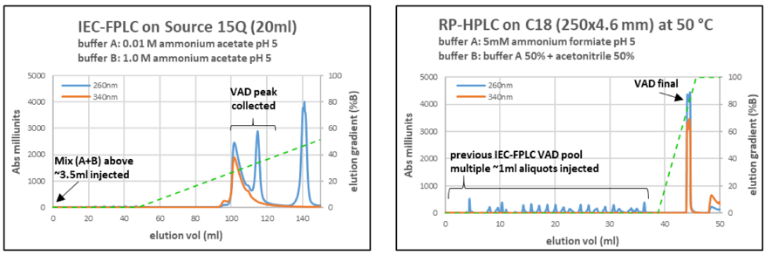


**Fig. S10. Related to Methods. VMN and VAD synthesis and purification.**

**(A)** Scheme of the reactions for VMN and VAD synthesis. **(B)** List of reagents used in mix (A) for VMN synthesis and mix (A+B) for VAD synthesis. **(C)** Representative IEC-FPLC purification of VMN from a typical mix (A). **(D)** Representative IEC-FPLC + RP-HPLC purification of VAD from a typical mix (A+B). Starting from vacor, typical conversion yields obtained via this protocol were 70% for VMN and 50% or less for VAD that required two chromatographic steps for purification.

**Table S1. X-ray data collection and structural refinement statistics.**

| | |
|---|---|
| **Data collection** | |
| Space group | P1 |
| a, b, c (Å) | 39.06, 51.01, 76.56 |
| α, β, γ (°) | 103.49, 101.78, 96.07 |
| Resolution (Å) | 46.60-1.69 (1.72-1.69) |
| $R_{merge}$ | 0.06 (0.96) |
| $R_{meas}$ | 0.07 (1.13) |
| $R_{pim}$ | 0.04 (0.59) |
| Mean I/σ (I) | 13.7 (1.8) |
| $CC_{1/2}$ | 1.00 (0.69) |
| Total reflections | 428,508 (18,156) |
| Unique reflections | 60,345 (2,695) |
| Completeness (%) | 96.6 (84.6) |
| Multiplicity | 7.1 (6.7) |
| **Refinement** | |
| $R_{work}$ | 0.19 |
| $R_{free}$ | 0.23 |
| RMS bonds (Å) | 0.00 |
| RMS angles (°) | 0.85 |
| Ramachandran favored (%) | 98.68 |
| Ramachandran outliers (%) | 0 |
| Rotamer outliers (%) | 0.19 |
| Clashscore | 2.15 |
| Average B-factor of the protein ($Å^2$) | 41.99 |
| Average B-factor of the ligand ($Å^2$) | 34.14 |
| C-beta outliers | 0 |

1. The statistics are based on the calculations from Aimless and MolProbity.

2. The numbers in parentheses represent the highest resolution shell.

3. $R_{merge} = \sum_{hkl} \sum_j |I_{hkl,j} - < I_{hkl} > | / (\sum_{hkl}\sum_j I_{hkl,j})$; $R_{meas} = \sum_{hkl} [N/(N-1)]^{1/2} \sum_j |I_{hkl,j} - < I_{hkl} > | / (\sum_{hkl}\sum_j I_{hkl,j})$; $R_{pim} = \sum_{hkl} [1/(N-1)]^{1/2} \sum_j |I_{hkl,j} - < I_{hkl} > | / (\sum_{hkl}\sum_j I_{hkl,j})$

4. $R_{work} = \sum_{hkl} |Fobs_{hkl} - Fcalc_{hkl}| / \sum|Fobs_{hkl}|$; $R_{free}$ is equivalent to $R_{work}$, with 5% of data excluded from refinement process. $|Fobs_{hkl}|$ and $|Fcalc_{hkl}|$ represent the observed and calculated structure factor amplitudes.
